# Transforming representations of movement from body- to world-centric space

**DOI:** 10.1101/2020.12.22.424001

**Authors:** Jenny Lu, Elena A. Westeinde, Lydia Hamburg, Paul M. Dawson, Cheng Lyu, Gaby Maimon, Shaul Druckmann, Rachel I. Wilson

**Affiliations:** Department of Neurobiology and Howard Hughes Medical Institute, Harvard Medical School, Boston, MA, USA; Department of Neurobiology, Stanford University, Stanford, CA, USA; Laboratory of Integrative Brain Function and Howard Hughes Medical Institute, The Rockefeller University, New York, NY, USA

## Abstract

When an animal moves through the world, its brain receives a stream of information about the body’s translational movement. These incoming movement signals, relayed from sensory organs or as copies of motor commands, are referenced relative to the body. Ultimately, such body-centric movement signals must be transformed into world-centric coordinates for navigation^1^. Here we show that this computation occurs in the fan-shaped body in the *Drosophila* brain. We identify two cell types in the fan-shaped body, PFNd and PFNv^2,3^, that conjunctively encode translational velocity signals and heading signals in walking flies. Specifically, PFNd and PFNv neurons form a Cartesian representation of body-centric translational velocity – acquired from premotor brain regions^4,5^ – that is layered onto a world-centric heading representation inherited from upstream compass neurons^6–8^. Then, we demonstrate that the next network layer, comprising hΔB neurons, is wired so as to transform the representation of translational velocity from body-centric to world-centric coordinates. We show that this transformation is predicted by a computational model derived directly from electron microscopy connectomic data^9^. The model illustrates the key role of a specific network motif, whereby the PFN neurons that synapse onto the same hΔB neuron have heading-tuning differences that offset the differences in their preferred body-centric directions of movement. By integrating a world-centric representation of travel velocity over time, it should be possible for the brain to form a working memory of the path traveled through the environment^10–12^.

---

Natural locomotor behavior combines body rotation with forward and lateral body translation. All these movements contribute to the path traveled by the animal through the world. For example, when a fly walks in a curved path to the left, it steps forward while rotating left and also sidestepping laterally to the left^13,14^ (Fig. 1a, Extended Data Fig. 1). The brain would underestimate the path’s curvature unless it tracked the body’s lateral movement in addition to tracking its forward movement.

**Figure 1.**
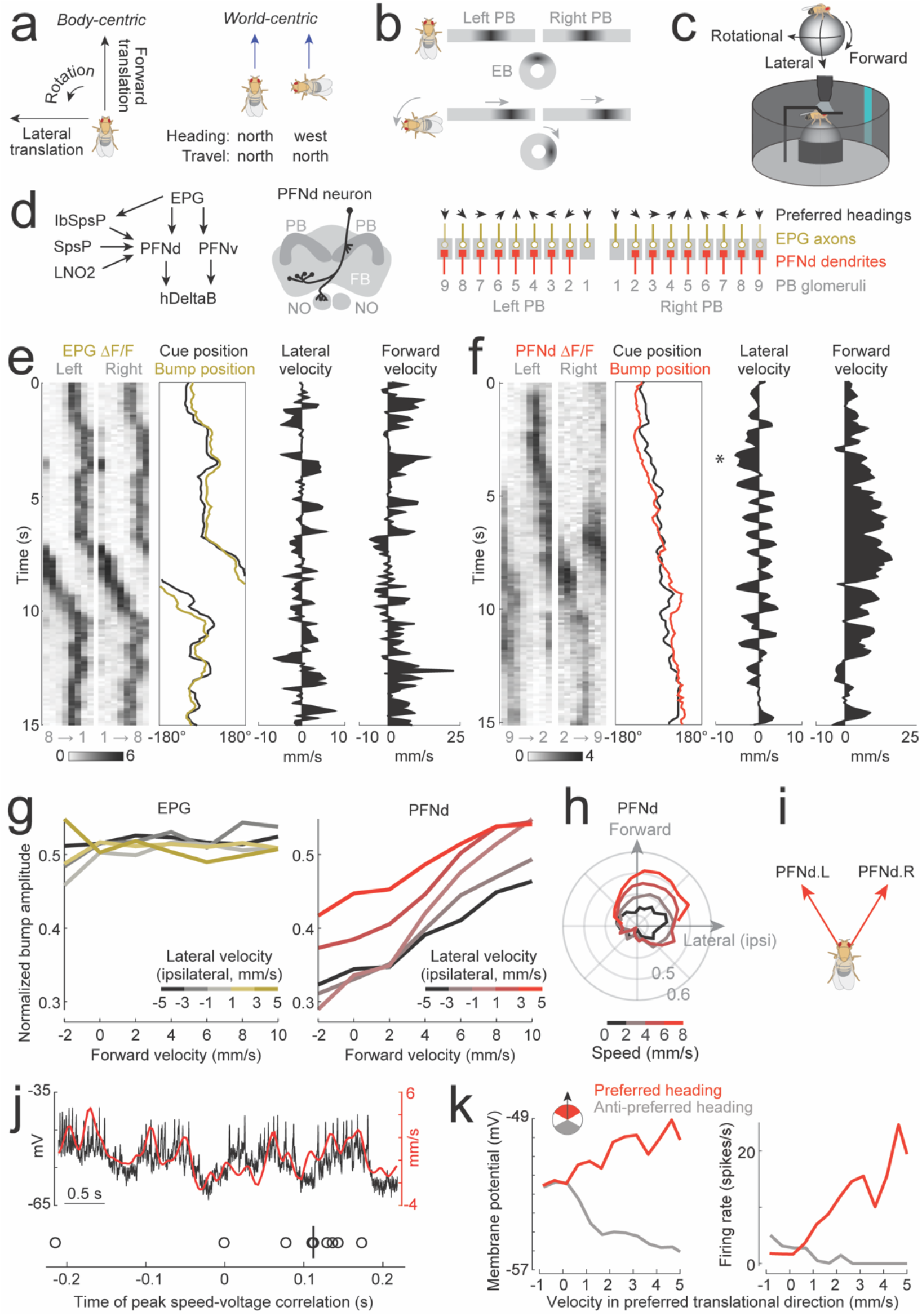
PFNd neurons are tuned to heading and translational velocity. a. Left: Movement axes in body-centric coordinates. Translation is defined as movement along the forward and lateral axes. Right: In world-centric coordinates, walking straight while heading north produces the same travel direction (blue arrow) as rightward lateral walking while heading west. b. EPG neuron dendrites form a topographic map of heading in the ellipsoid body (EB) while their axons form two linearized maps in the protocerebral bridge (PB). When the fly rotates clockwise, the bump of activity in the EB rotates counter-clockwise, and PB bumps move leftward. Here and elsewhere, we depict brain anatomy from the posterior side of the head. c. Two-photon calcium imaging as a fly walks on a spherical treadmill in closed loop with a visual cue. d. Left: cell types in this study and their connections, based on the partial brain connectome^9^. We include here all the major unilateral inputs to PFNd; bilateral PFNd inputs are omitted for clarity (Extended Data Fig. 3). Middle: morphology of a single PFNd neuron^2^. Synaptic inputs relevant to this study arrive via the cell’s dendrites in the PB and nodulus (NO), while synaptic outputs are in fan-shaped body (FB). Right: PFNd dendrites occupy PB glomeruli 2-9. EPG axons target glomeruli 1-9, but the line we used to drive expression of the calcium indicator omits the EPG neurons in the most lateral glomeruli (shading). Arrows show relative heading preferences of EPG neurons. e. EPG population activity in the PB as a fly walks in closed loop with a visual cue. f. Same for the PFNd population. When lateral velocity is mainly leftward (*), the left bump amplitude is larger than the right. g. Left: Normalized EPG bump amplitude versus forward velocity. Data are binned by lateral velocity in the ipsilateral direction (i.e., right for EPG projections to the right PB and left for projections to the left PB). Data from the right and left PB are combined and then averaged across flies (n = 5 flies). Right: same for PFNd neurons in the PB (n = 16 flies). h. Normalized PFNd bump amplitude in the PB, versus body-centric translational direction. Data are binned by speed. Lateral velocity is expressed in the direction ipsilateral to the imaged PB, allowing us to combine data from the right and left PB before averaging across flies (n = 16 flies). i. Preferred body-centric translational directions of the PFNd.R and PFNd.L populations. Based on a linear fit to the data in (g), we estimate that these neurons prefer translation directions of ±31° (see Methods and Extended Data Fig. 2). j. Top: membrane potential of a PFNd neuron, overlaid with the fly’s walking speed in this neuron’s preferred body-centric translational direction. Bottom: lag between velocity (in the cell’s preferred body-centric translational direction) and voltage. Each symbol is a fly (n=9 cells in 7 flies, vertical bar shows median lag = 0.113 s). k. Left: mean membrane potential of an example PFNd cell, plotted versus velocity in this neuron’s preferred body-centric translational direction. Data are divided into three bins based on the proximity of the fly’s heading to the neuron’s preferred heading (black arrow). Right: same for spike rate.

A walking insect can measure its forward and lateral velocity using proprioceptive signals and copies of internal motor command signals^15^, as well as optic flow signals^16,17^. All these velocity signals arrive in body-centric coordinates. Ultimately, however, if the brain is to construct a memory of the fly’s navigation path^10,18^, it must convert these body-centric signals into world-centric coordinates^1,19^. For example, moving forward while facing north should be represented by the brain’s navigation systems as equivalent to moving right while facing west (Fig. 1a) – even if these two situations involve very different peripheral sensory signals and motor commands.

If this computation is performed in the fly brain, it would likely occur downstream from the ellipsoid body (EB) of the central complex. The EB contains a topographic map of heading which is anchored to multimodal environmental cues^6,20,21^ – in other words, a world-centric heading representation. EB projection neurons (EPG neurons or “compass neurons”) send this heading information to the protocerebral bridge (PB)^7,8^ (Fig. 1b). Indeed, a recent study in the honeybee showed that, in some regions of the central complex downstream from the PB, there are incoming projections from neurons that encode the direction of optic flow during flight^22^. This result suggests a possible fusion of translational velocity signals and heading signals in the central complex. To pursue this idea, we turned to the fly *Drosophila melanogaster*, where we can use transgenic driver lines^2,3^ to target central complex neurons with well-defined synaptic connectivity^19^.

## Neurons encoding heading and velocity

To determine whether heading information is integrated with velocity information downstream from EPG neurons, we used specific driver lines^3^ to express a fast calcium indicator (jGCaMP7f)^23^ in various cell types in the PB. We imaged these neurons using two-photon excitation microscopy as the fly walked freely on a spherical treadmill, surrounded by a visual virtual reality environment^24^ (Fig. 1c). This environment was a 360° panorama displaying a prominent heading cue – namely, a bright object which rotated rightward when the fly made a fictive leftward turn, and *vice versa*.

We focused on a specific PB cell type (PFNd) downstream from compass neurons (Fig. 1d), because PFN neurons are anatomically well-positioned to receive both heading information and translational velocity information^22^. We found that PFNd dendrites form two complete maps of heading, one on each side of the PB (PFNd.L and PFNd.R, Fig. 1d), resembling the two complete maps in EPG axons (Fig. 1e,f). As the fly rotates clockwise, there is a “bump” of activity which moves leftward across the PFNd.R population, and another bump that moves leftward across the PFNd.L population. Overall, the position of the bump is correlated with heading, as in EPG neurons (Extended Data Fig. 2).

Moreover, we found that PFNd neurons also show direction-selective responses to translational movement. Specifically, the PFNd.R bump amplitude increases when the fly translates forward and right, whereas PFNd.L bump amplitude increases when the fly translates forward and left (Fig. 1g-i, Extended Data Fig. 2). Thus, PFNd neurons are sensitive to both forward and lateral velocity. These neurons are somewhat correlated with rotation as well (Extended Data Fig. 2), but this is unsurprising because lateral and rotational velocity are highly correlated during walking (Extended Data Fig. 1)^13,25^; moreover, as we will see, the function of these neurons is most relevant to translational movement.

We estimated the preferred translational direction of each population by fitting a linear function to their two-dimensional tuning profile (Extended Data Fig. 2); this analysis suggested that PFNd.R and PFNd.L neurons prefer translation at an angle of 31° and −31°, respectively, relative to heading (Fig. 1i). Graded velocity increases in the preferred direction produce graded increases in bump amplitude (Extended Data Fig. 2). By contrast, EPG neurons themselves are relatively insensitive to translational velocity (Fig. 1g).

Next, we used whole-cell recordings to measure the relative latency of PFNd activity and behavior. When we cross-correlated voltage and velocity in the cell’s preferred translational velocity direction, we found that voltage typically lagged velocity by about 0.1 s (Fig. 1j). This lag implies that PFNd neurons are not directly causal for these changes in velocity. In other words, PFNd neurons are measuring velocity rather than commanding velocity.

These electrophysiological recordings also revealed that effect of velocity (in the cell’s preferred body-centric direction) is to multiplicatively scale the effect of heading. For a given heading, the cell’s membrane potential depends quite linearly on velocity (Fig. 1k). High velocities amplify the depolarizing effect of the preferred heading, as well as the hyperpolarizing effect of the anti-preferred heading (Fig. 1k).

In summary, PFNd neurons support a conjunctive code for several navigational variables. The spatial phase of the activity bumps (i.e. the position of the bump along the PB) encodes heading, which is a world-centric variable. Meanwhile, the amplitude of each bump scales linearly with translational velocity in a specific direction, with the right and left PFNd populations tuned to different body-centric directions.

## Velocity coding via graded release from inhibition

To understand the origin of the conjunctive code in PFNd neurons, we identified the inputs to PFNd neurons in the hemibrain connectome^9^ (Extended Data Fig. 3). We then used specific driver lines to express jGCaMP7f in the cell types that provide major unilateral input to PFNd (Fig. 1d, Extended Data Fig. 3) to understand how the left and right PFNd populations acquire their translational direction selectivity. We found strong direction-selective translational velocity signals in two cell types, SpsP and LNO2. These two cell types are output projection neurons from little-studied brain regions that have been implicated as premotor regions associated with locomotion (the SPS and LAL, respectively)^4,5^.

Notably, we found that SpsP and LNO2 neurons are anti-correlated with forward velocity (Fig. 2a-f), rather than being positively correlated as PFNd neurons are. To determine whether SpsP and LNO2 neurons might be inhibitory, we reconstructed examples of these neurons in a second large-scale EM dataset^26^ which allows us to leverage machine learning to predict the neurotransmitters associated with each cell^27^. We found that SpsP and LNO2 output synapses were mainly scored as glutamatergic. Glutamate is often an inhibitory neurotransmitter in the *Drosophila* brain^28^, and indeed we confirmed that optogenetic activation of SpsP neurons produces PFNd neuron hyperpolarization, with the characteristic pharmacological signature of monosynaptic inhibition via glutamate-gated chloride (GluCl) channels (Fig. 2g). We also confirmed that a split-Gal4 hemidriver that reports vesicular glutamate transporter expression^29^ can drive expression in LNO2 neurons (Extended Data Fig. 4). Thus, both types of premotor projection neurons are likely inhibitory. This means that, as forward velocity increases, PFNd neurons receive graded disinhibition from premotor regions.

**Figure 2.**
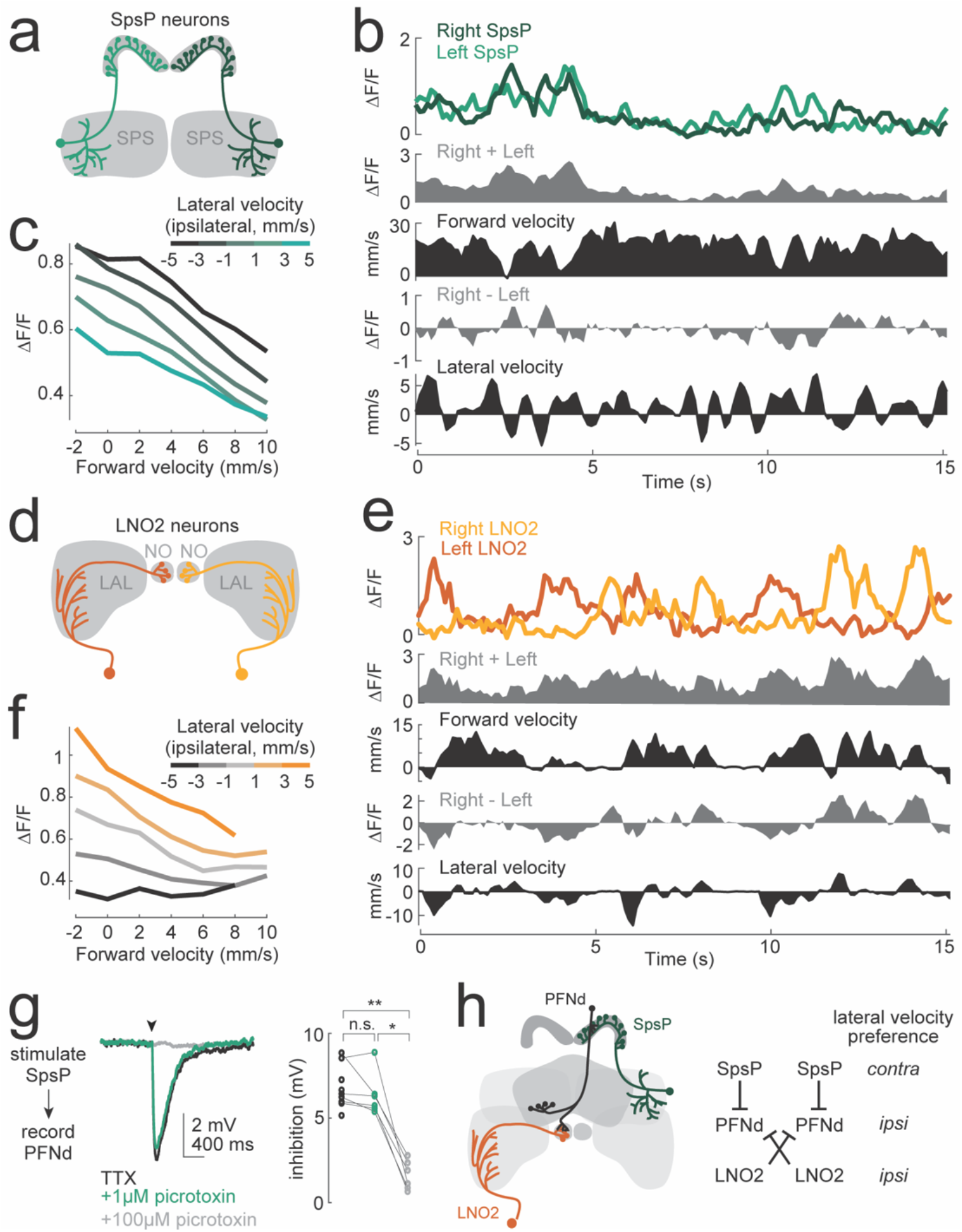
Velocity tuning in PFNd neurons from graded release of inhibition. a. Each SpsP neuron receives input from the superior posterior slope (SPS) and synapses onto PFNd dendrites across the ipsilateral PB. b. SpsP activity as a fly walks in closed loop with a visual cue. c. SpsP activity versus forward velocity. Data are binned by lateral velocity in the ipsilateral direction and averaged across flies (n = 8 flies). d. Each LNO2 neuron receives input from the lateral accessory lobe (LAL) and synapses onto all the PFNd neurons in the ipsilateral NO. e. LNO2 activity as a fly walks in closed loop with a visual cue. f. LNO2 activity versus forward velocity. Data are binned by lateral velocity in the ipsilateral direction and averaged across flies (n = 4 flies). g. Left, whole-cell voltage response of a PFNd neuron to optogenetic stimulation of SpsP neurons (arrowhead), recorded in TTX (to isolate monosynaptic input), in TTX + 1 μM picrotoxin (to block GABA_A_ receptors^51^), and TTX + 100 μM picrotoxin (to block GluCl receptors^28^). Each trace is an average of >50 trials. Right, amplitude of stimulus-evoked inhibition. Within each condition, each symbol is a cell (n=9 flies, with picrotoxin for 6 of these flies, ** P=2.67×10^−4^ and * P=7.02×10^−4^, paired-sample t-tests with Bonferroni-corrected α = 0.0167). h. Summary of lateral velocity preferences. Because PFNd dendrites in the NO are contralateral to their dendrites in the PB, the right LNO2 neuron synapses onto PFNd.left, and vice versa.

Recall that PFNd neurons are sensitive to lateral velocity. We found that SpsP and LNO2 neurons are also sensitive to lateral velocity (Fig. 2a-f, Extended Data Fig. 5); moreover, their lateral direction selectivity can explain the observed lateral direction selectivity of PFNd neurons (Fig. 2h). Specifically, when the fly walks to the right, SpsP and LNO2 neurons inhibit PFNd.L, while also disinhibiting PFNd.R. Thus, these premotor projection neurons can account for both the lateral and forward velocity tuning of PFNd neurons.

PFNd neurons also receive a major input from IbSpsP neurons (Extended Data Fig. 5). However, we found that IbSpsP neurons are not very selective for the direction of translational motion (Extended Data Fig. 5). Thus, they are unlikely to account for direction-selective velocity tuning in PFNd neurons.

To summarize, PFNd neurons in each brain hemisphere inherit a map of heading direction from the compass system. These maps are under the inhibitory control of premotor projection neurons which encode both the speed and direction of body movement. Lateral movement releases the ipsilateral PFNd heading map from inhibition, while increasing inhibition of the contralateral map. By contrast, forward movement disinhibits both maps. In this way, body-centric velocity information is layered onto the world-centric map of heading.

## A complete Cartesian system for velocity

Next, we investigated another cell type, PFNv, with a morphology nearly identical to PFNd neurons (Fig. 3a). Like PFNd neurons, PFNv neurons are downstream from compass neurons (Fig. 1d, Extended Data Fig. 3). Accordingly, we found that PFNv neurons also form a heading map in each half of the PB (Fig. 3b), with a bump of activity whose position correlates with heading (Extended Data Figure 6).

**Figure 3.**
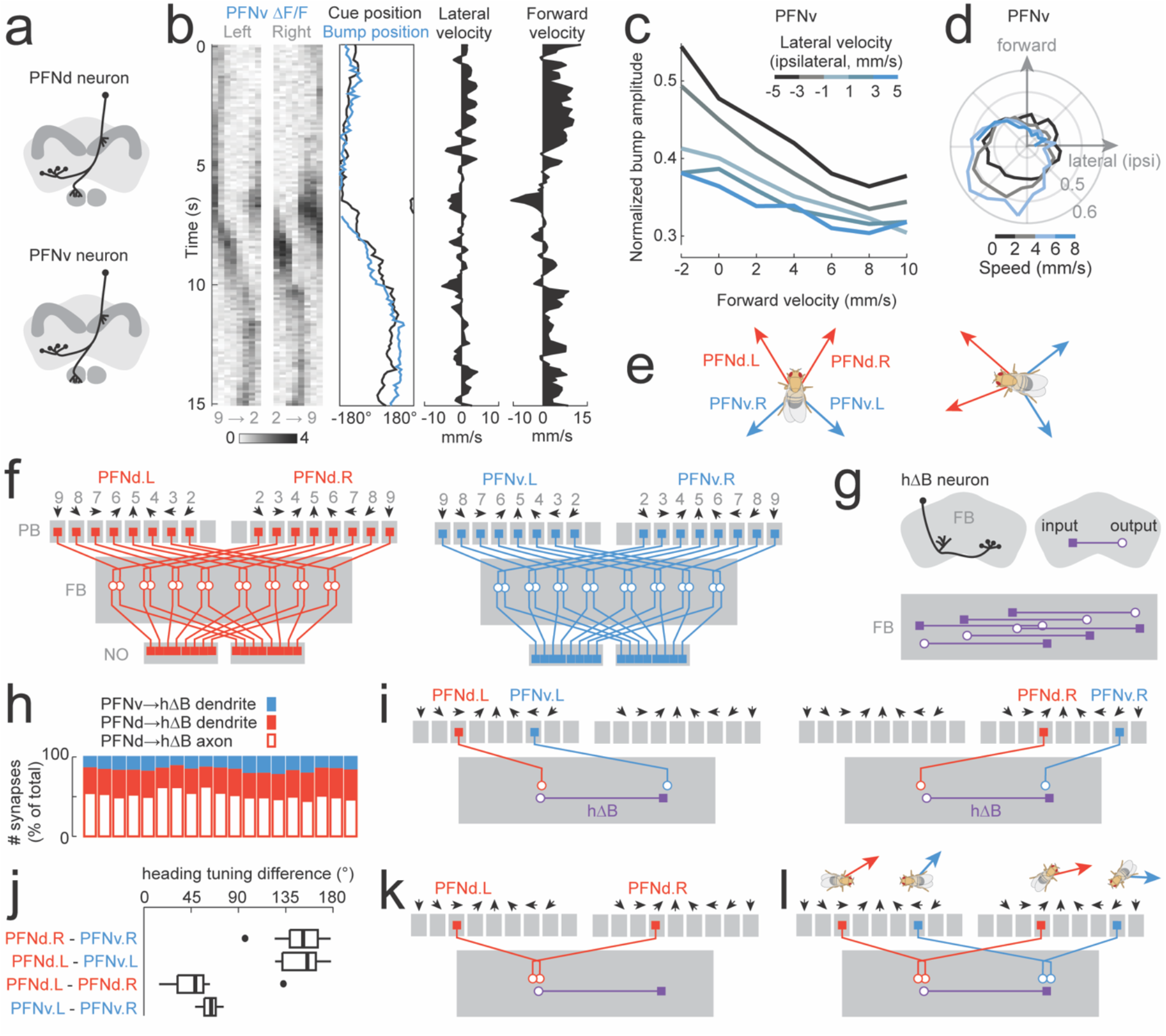
PFNd and PFNv neurons form a Cartesian coordinate system for translational velocity. a. PFNd and PFNv neurons have similar morphologies, except their dendrites reside in the dorsal and ventral NO2, respectively^2^. b. PFNv population activity in the PB as a fly walks in closed loop with a visual cue. c. Same as Fig. 1g but for PFNv (n = 11 flies). d. Same as Fig. 1h but for PFNv (n = 11 flies). e. Left: The full PFNd/v coordinate system. Based on a linear fit to the data in (c), we estimate that PFNv.L neurons prefer a translation direction of +132°, whereas PFNv.R neurons prefer −132°. (see Methods). Note that PFNv neurons prefer contralateral movement, whereas PFNd neurons prefer ipsilateral movement. Right: when the fly rotates, the PFNd and PFNv coordinate system rotates accordingly. f. Schematized projections of the PFNd and PFNv populations, from the hemibrain connectome^9^. Note that the mapping from PB glomeruli to FB horizontal locations is the same for PFNd and PFNv. For each cell type, each half of the PB contains a complete heading map (arrows) which is projected onto the full horizontal axis of the FB. g. Top: Morphology of an hΔB neuron. This cell type has a dendrite in the FB and an axon that projects halfway across the horizontal extent of the FB (schematized as a filled square and open circle). Bottom: hΔB neurons tile the horizontal extent of the FB (schematized as a rectangle). All hΔB neurons reside in the same FB layer but are vertically offset for clarity in this schematic. h. Synapses onto hΔB dendrites and axons from PFNv and PFNd neurons. Each vertical bar represents one hΔB neuron (n = 19 neurons). i. Schematized projections onto a single hΔB neuron from each side of the PB. PFNd neurons target both hΔB neuron axons and dendrites, but their dominant input is axonal. For a given side of the PB, PFNd and PFNv inputs have different heading tuning. j. Analysis of hemibrain connectome data. Each PFNd and PFNv neuron in the connectome was assigned a preferred heading, and each PFN→hΔB connection was weighted by the number of synapses in that connection. Then, for each hΔB neuron, we computed the weighted average of the preferred headings of its PFN inputs; we did this separately for each input population (PFNd.R, PFNv.R, PFNd.L, PFNv.L) onto each hΔB neuron. Finally, for each hΔB neuorn, we computed the absolute pairwise difference between these averages. Each boxplot summarizes results for all 19 hΔB neurons; thick line = median, box = interquartile range, whiskers = full range omitting one outlier (•). k. Schematized projections onto a single hΔB neuron from PFNd neurons. Note that PFNd.L and PFNd.R inputs to this neuron have different heading tuning. The same is true for PFNv.L and PFNv.R inputs. l. A schematic hΔB neuron and its top PFN inputs. Above each PFN neuron is a fly with a body-centric translational velocity vector (red or blue) denoting the preference of that PFN neuron. The rotation of the fly denotes the preferred heading of the PFN neuron. Note that all four PFN neurons have similar world-centric travel direction preferences.

Notably, we found that PFNv bump amplitude encodes translational velocity along directions that are opposite to those encoded by PFNd neurons. Whereas PFNd neurons are excited by forward and ipsilateral velocity, PFNv neurons prefer backward and contralateral velocity (Fig. 3c-e).

Because PFNd and PFNv neurons have such different translational direction selectivity, we might expect them to receive different inputs from the brain’s premotor centers (SPS and LAL^4,5^). The hemibrain connectome^9^ confirms this prediction: SpsP and LNO2 neurons make many synapses onto PFNd neurons, but they make almost no synapses onto PFNv neurons. Rather, PFNv neurons are downstream from other LAL projection neuron types (LNO1 and LNO3; Extended Data Fig. 3) which contribute relatively little to PFNd input. LNO1 neuron activity was not strongly correlated with body velocity under our experimental conditions (Extended Data Fig. 7), so it seems more likely that LNO3 neurons are the major drivers of PFNv responses, although we were not able to obtain a selective driver for LNO3 neurons that would allow us to test this idea.

In summary, PFNd and PFNv neurons together constitute a complete set of Cartesian axes for encoding translational velocity (Fig. 3e). This representation is in body-centric coordinates. Thus, when the fly rotates, the PFN velocity-vector representation rotates along with the fly’s heading (Fig. 3e). By conjointly encoding the full 360° of body-centric translational velocity directions and the full 360° of heading directions, PFNd and PFNv neurons are poised to transform velocity representations from a body-centric coordinate system to a world-centric coordinate system.

## Integrating opponent populations

Next, we examined what happens downstream from PFNd and PFNv neurons. Both of these cell types project to the fan-shaped body (FB), where their axonal projections preserve the rough topography of the heading map^2^. The hemibrain connectome^9^ reveals that PFNd and PFNv projections from the same PB glomerulus are precisely overlaid in the FB, and they converge onto the same cell type, called hΔB (Fig. 3g).

This projection pattern seems puzzling. PFNd and PFNv neurons from the same PB glomerulus have opposing velocity preferences (Fig. 3e). Thus, the convergence of these projections would seem to cancel the directional information provided by each PFN population.

The resolution to this puzzle lies in the subcellular targeting of these synaptic connections. Each hΔB neuron spans half of the horizontal extent of the FB, with a dendrite at one pole and axon terminals at the other (Fig. 3g). In analyzing the connectome, we discovered that PFNv neurons synapse selectively onto hΔB dendrites, whereas PFNd neurons synapse onto both hΔB axon terminals and dendrites, with a preference for axon terminals (Fig. 3h).

If we take the perspective of an individual hΔB neuron, we see that its PFNd and PFNv inputs from the left PB will have opposite heading tuning (Fig. 3i, left). Similarly, PFNd and PFNv inputs from the right PB will also have opposite heading tuning (Fig. 3i, right). If PFNd neurons exclusively targeted hΔB axon terminals (as schematized in Fig. 3i), then the heading difference would be exactly 180°, but because PFNd neurons also target hΔB dendrites, the heading difference is closer to 150° (Fig. 3j). Recall that PFNd and PFNv neurons from the PB on the same hemisphere have nearly opposite preferred directions of translational velocity, which roughly matches this heading difference (Fig. 3e).

To summarize, PFNd and PFNv inputs to the same hΔB neuron have nearly opposite heading tuning, and also nearly opposite translational velocity tuning. As a result, their translational velocity signals should combine constructively, rather than destructively. The key point is that PFNd and PFNv neurons that wire together (onto the same hΔB neuron) have heading-tuning differences that offset their translational direction tuning differences. This is a notable example of a geometrical computation in the brain which relies on the precise geometry of subcellular synaptic targeting.

## Interhemispheric integration

There is yet another important feature of the PFN projection pattern: projections from the right and left copies of the PB heading map are not aligned in the FB^2^. Rather, there is a systematic shift, so that projections from the right PB are shifted leftward, while projections from the left PB are shifted rightward (Fig. 3k). Thus, when we analyzed the hemibrain connectome from the perspective of an individual hΔB neuron, we found that PFNd inputs from the right and left have preferred headings that differ by ~45°, on average (Fig. 3j). Note that this heading-tuning difference is opposite to the translational direction tuning difference in the two PFNd populations (Fig. 3k). This matched shift allows the hΔB neuron to constructively combine information from the left and right hemispheres.

The same matched shift occurs for PFNv neurons, although here there is a somewhat larger heading tuning difference between left and right inputs to an individual hΔB neuron (Fig. 3j). Accordingly, there is also a somewhat larger difference in the preferred translational directions of right and left PFNv populations (as compared with the difference in the right-left PFNd populations, Fig. 3e). Again, the key point is that the PFN neurons that wire together have heading-tuning differences that offset their translational direction tuning differences.

## A body-to-world coordinate transformation

To see the functional consequence of this anatomy, consider again the perspective of an individual hΔB neuron (Fig. 3l). Its inputs have diverse preferred headings, and diverse body-centric direction preferences. What these inputs have in common is their preferred world-centric travel direction (Fig. 3l). Thus, the identity of the active hΔB neurons should encode the direction of world-centric travel. Travel in a given direction should activate the same hΔB neurons, regardless of whether the fly is walking right or left, forwards or backwards. In other words, the brain’s translational velocity-vector should be transformed from body-centric coordinates to world-centric coordinates.

To investigate this idea, we implemented a computational model of the network, comprising 40 PFNd, 20 PFNv, and 19 hΔB neurons, identical to the cell numbers in the hemibrain connectome^9^. For simplicity, we directly modeled the activity of PFN neurons as a function of heading and body-centric translational velocity, rather than modeling the network inputs to PFN neurons. In the model, each PFN neuron is cosine-tuned to heading. The amplitude of its heading response is multiplicatively scaled by the nonnegative component of the fly’s translational velocity in that cell’s preferred body-centric direction (Fig. 4a). This follows what we see in PFN membrane voltage data, where velocity multiplicatively scales the cell’s heading tuning (Fig. 1k). PFN→hΔB connectivity is taken directly from the connectome, with weights proportional to the number of synapses per connection (Fig. 4b). This direct approach is a departure from previous studies where connections weights from connectome data were computationally optimized to achieve a particular modeling result^30,31^; here, connections were simply taken directly from data, with no optimization. Finally, each hΔB neuron in the model simply sums its PFN inputs.

**Figure 4.**
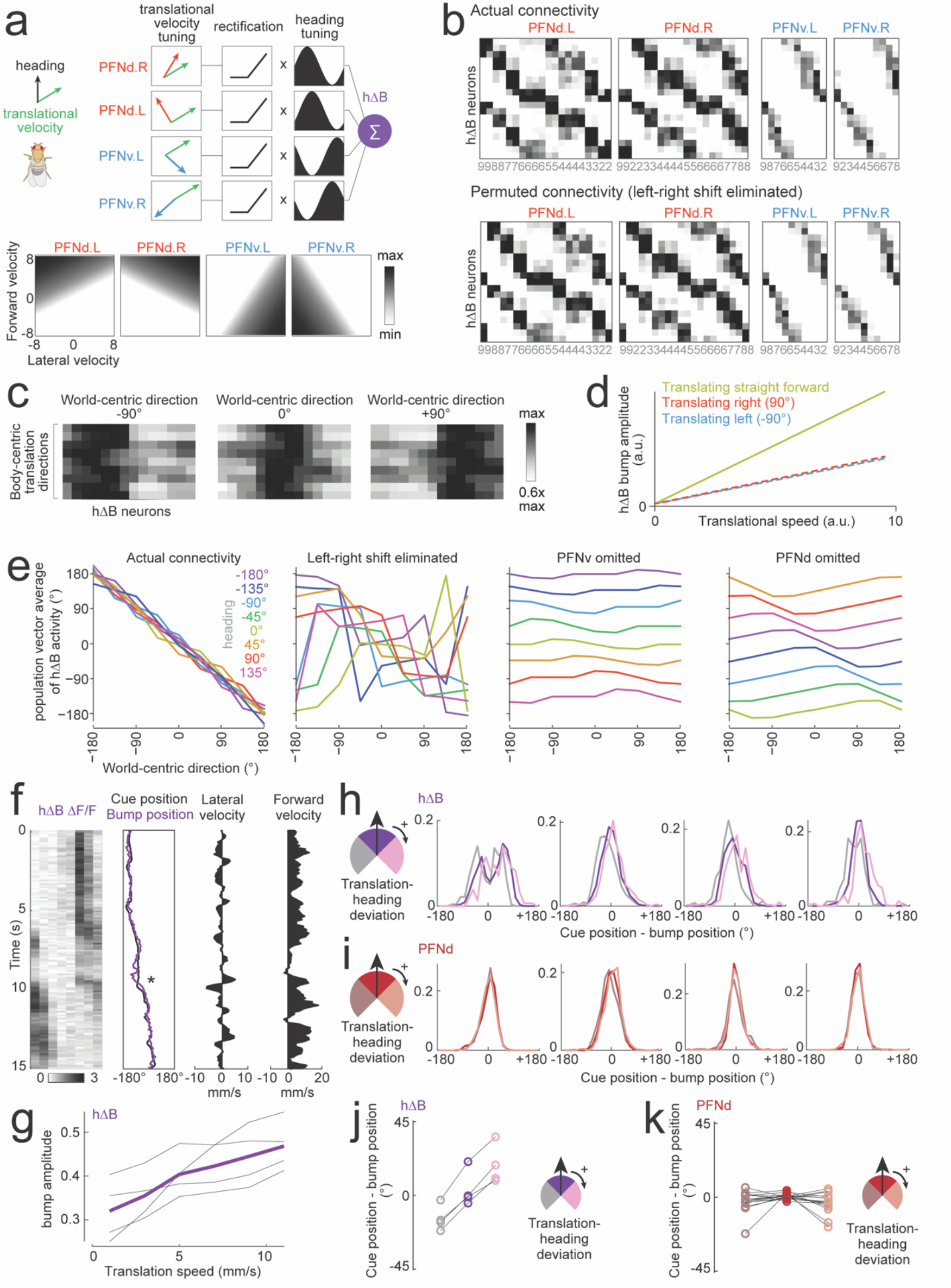
From body-centric velocity and heading to world-centric velocity. a. In the model, 40 PFNd neurons and 20 PFNv neurons converge onto 19 hΔB neurons. Each PFN population is tuned to translational velocity as shown in the grayscale heatmaps below. The fly’s translational velocity (green arrow) is projected onto each cell’s preferred translational direction (red/blue arrows); the result is then rectified and used to scale the cell’s heading response. Each PFN neuron is cosine-tuned to heading, with a heading response in the range [0,1], and a preferred heading dictated by its cognate PB glomerulus (Fig. 3f). These PFN firing rates are then weighted by the PFN→hΔB connection matrix from the hemibrain connectome and summed in hΔB neurons. Finally, independent Gaussian noise is added to each hΔB neuron to emulate the noisiness of real networks. b. Top: the PFN→hΔB connection matrix from the hemibrain connectome, with darker colors representing more synapses, and thus stronger weights in the model. Rows are hΔB neurons, columns are PFN neurons. Within each PFN population, columns are sorted by PFN heading tuning, as inferred from the PFN neuron’s PB glomerulus^2^. Gray numbers denote PB glomeruli. Recall that PFNd neurons target both hΔB axons and hΔB dendrites, and so there are two “hotspots” per PFNd column. c. Model: population activity of hΔB neurons. The fly’s path is a straight line at a fixed speed. In consecutive rows within the same block, heading is rotated clockwise, while body-centric velocity is rotated counterclockwise by the same angle, so that the world-centric travel vector is the same for all rows in a block. Activity is normalized to the maximum in each row. Note that the position of the hΔB bump is essentially unchanged within a block, for all three blocks (three world-centric travel directions). d. Model: hΔB bump amplitude versus translational speed, for three heading directions. e. Model: position of the hΔB bump (computed as the population vector average or PVA) as a function of world-centric travel direction. Results are shown for 7 different headings. In the full model, PVA is proportional to travel direction and invariant to heading. In a model with permuted connectivity that removes the left-right phase shift (shown in b), world-centric direction encoding is disrupted, but not completely eliminated, due to the fact that PFNd and PFNv connections are still properly arranged relative to each other. When either PFNv or PFNd neurons are omitted, world-centric direction encoding is almost completely eliminated. f. Data: hΔB population activity in the FB as a fly walks in closed loop with a visual cue. The FB is segmented into vertical columns, and ΔF/F is measured in each column. When lateral speed is high relative to forward speed, the position of the hΔB bump deviates from the fly’s heading (asterisk). g. Data: hΔB bump amplitude (normalized to the maximum in each fly) versus translational speed. Each gray line is a fly; purple is the mean across flies. h. Histograms showing the difference between the cue position and the hΔB bump position. These values are mean-centered within experiment, and then binned according to the deviation between the fly’s translational direction and its heading. Results are shown for four flies. When the fly’s translation direction deviates from its heading, the hΔB bump shifts away from the cue position. i. Same but for PFNd neurons. Here, the bump does not shift away from the cue position. j. Mean difference between the cue position and the hΔB bump position. Each set of connected symbols is one experiment (n=4 flies).The purple and gray distributions are significantly different, as are the purple and pink distributions (P=0.0030 and P=0.0038, respectively; paired-sample t-tests with Bonferroni-corrected α = 0.0167,). k. Same for PFNd (n=16 flies). The red distributions are not significantly different from the gray or peach distributions (P=0.0909 and P=0.0446, paired-sample t-test with Bonferroni-corrected α = 0.0167).

Notably, in this model, the position of the hΔB activity bump tracks the fly’s world-centric travel direction (Fig. 4c) and the amplitude of the bump scales with travel speed, although the gain of this relationship is highest when the fly is walking straight forward (Fig. 4d). The model’s travel direction encoding is remarkably good: for a given world-centric travel direction, the position of the hΔB bump is essentially always the same, regardless of whether the fly’s heading direction matches it travel direction (Fig. 4e). Travel direction encoding is disrupted if we permute the connectivity matrix to remove the 45° phase shift in hΔB inputs from left and right PFN populations (Fig. 4b,e). It is even more disrupted if we remove either PFNv or PFNd neurons from the model (Fig. 4e). Although PFNv neurons contribute only a relatively small number of synapses (Fig. 3h), they are essential because their velocity tuning opposes that of PFNd neurons. Unless both opponent populations are functional, they cannot push the hΔB bump to the correct position.

Finally, we tested the predictions of this model by imaging jGCaMP7f in hΔB neurons. Although hΔB dendrites and axons are intercalated, calcium signals should be dominated by axons, given the concentration of calcium channels at presynaptic terminals. Thus, we would expect to see a moving bump of activity whose amplitude scales with travel speed. This is indeed what we observed (Fig. 4f,g). The bump position generally followed the visual heading cue, which is expected whenever the fly is walking fairly straight. Notably, however, the hΔB bump shifted away from the cue when the fly’s lateral speed was high relative to its forward velocity (Fig. 4f) – i.e., when travel direction deviated from heading (Fig. 4h). The observed bump shift was relatively small, which is what we would expect, given the limited temporal resolution of calcium imaging and the fact that travel-heading deviations are very transient during normal walking (Fig. 4f). Importantly, however, we saw a quantitatively similar shift in every hΔB experiment, and we did not observe this type of shift in PFNd experiments (Fig. 4i-k, Extended Data Fig. 8) or in PFNv experiments (Extended Data Fig. 8).

In summary, while the PFN bump position faithfully tracks the fly’s heading direction, the hΔB bump position is sensitive to the intermittent deviations between travel and heading that occur during normal walking. This result supports the idea that hΔB neurons form a map of travel direction, rather than heading direction. Meanwhile, the amplitude of hΔB activity encodes travel speed. Together, these results imply that hΔB neurons constitute a vector-velocity representation of travel direction and speed in world-centric coordinates.

## Discussion

The transformation of velocity signals into world-centric coordinates is a prerequisite for computing a world-centric path during navigation. *Drosophila* and other insects can use path integration to navigate back to a familiar site in the absence of spatial position cues^10–12^. Path integration requires the brain to measure not only direction but also distance (or speed). A simple model proposed twenty-five years ago by Wittmann and Schwegler^32^ proposed that the compass system in the insect brain^33^ is the source of direction information; forward speed then multiplicatively scales the output of the compass system, and this representation is integrated over time to produce a vectorial representation of the distance traveled in each heading direction. A limitation of this model is that it assumes all translational velocity is straight ahead, with no lateral component.

This problem is solved in a model recently proposed by Stone, Webb, and colleagues^22^. Inspired by experimental work in honeybees, this model begins with a Cartesian coordinate system for translational velocity – specifically, one cell tuned to forward-right velocity, and another cell tuned to forward left velocity. In the model, each of these two neurons projects to a separate downstream “integrator” population which also receives a complete heading map. Heading signals are subtracted from translational velocity signals, and the result is summed over time in each of these two integrator populations. Note that, in the Stone-Webb model, there is no explicit representation of world-centric travel velocity. Rather, this model separately stores path components along two orthogonal axes of translation. It was predicted that PFN neurons (called CPU4 in bees) correspond to the integrator cells of the model. A limitation of this model is that it only works when there is no backward component to the agent’s movement.

Here we show that PFN neurons do indeed combine heading and translational velocity signals. However, we show there are actually four populations of PFN neurons that collectively tile the full 360° of velocity space in a Cartesian coordinate system. Contrary to the prevailing model, we found no evidence that PFNd or PFNv neurons integrate velocity information over time to yield a distance calculation.

Next, we show that this network is wired to perform a computation which has not previously been attributed to the insect brain: it computes an explicit representation of the body’s velocity in world-centric coordinates. This computation occurs at the next layer of the network, in hΔB neurons. The identity of the active neurons in the hΔB population represents world-centric travel direction. This means that, when the agent rotates, the hΔB coordinate system remains stable rather than rotating. Meanwhile, the amplitude of hΔB activity covaries with travel speed. A neural representation of speed is important to accurate navigation, and indeed there is behavioral evidence that path integration in walking^16,17^ and flying^34^ insects is sensitive to speed cues.

Moreover, our model shows that the crucial element in this computation is the precise targeting of PFNd and PFNv output synapses. In this wiring pattern, PFN neurons converge onto the same hΔB neuron if the difference in their translational-direction preference is offset by a shift in their heading-direction-preference. We suggest that an analogous wiring pattern might be found in the vertebrate brain, in the arrangement of inputs to world-centric velocity-vector cells^35^.

More generally, our work provides the first detailed mechanistic description of a body-to-world vectorial coordinate transformation in any species. This is potentially relevant to the many different vectorial codes in mammalian navigation systems^1^. Some of these vectorial codes operate in body-centric coordinates^35–38^, whereas others operate in world-centric coordinates^35,39–42^. It has been proposed that the outputs of body-centric vector cells are combined to produce world-centric vector cells in downstream network layers^43,44^. This would seem to be a crucial element of the neurobiology of navigation. Our results show that this does in fact occur – and indeed how it occurs – in an insect brain. A parallel study reaches many of the same conclusions^45^.

Ultimately, path position representations must be compared to internal spatial goals. Then, they must be transposed back into a body-centric reference frame for steering control^25^. By identifying wiring patterns in the connectome^19^, exploring these patterns in computational models, and testing these models through physiology experiments, it should be possible to achieve an understanding of all these computations at an algorithmic and biophysical level.

## Author contributions

Experiments were designed by J.L. and R.I.W. with input from E.W. and S.D. J.L. performed imaging experiments and analyses. E.W. performed electrophysiological experiments and analyses. L.H. and S.D. designed and implemented the computational model with input from J.L. and R.I.W. P.D. performed MCFO experiments. C.L. and G.M. provided the hΔB split-Gal4 line prior to publication.

## Acknowledgements

This study benefited enormously from the public release of the hemibrain connectome in January 2020 by the FlyEM Team at Janelia. Isabel Haber and Anna Li annotated neurons and synapses in the full adult female brain dataset (FAFB)^26^, and Nils Eckstein and Alexander Bates generated neurotransmitter predictions based on those data, using algorithms they designed and implemented with Jan Funke and Greg Jefferis. Will Dickson and Michael Dickinson shared modified FicTrac software, panels hardware support, and foam sphere machining help. Tanya Wolff, Gerry Rubin, and Vivek Jayaraman provided fly stocks. We thank Michael Dickinson and Amir Behbahani for helpful discussions, and members of the Wilson lab for comments on the manuscript. This work was supported by the Harvard Medical School Neurobiology Imaging Facility (NINDS P30 #NS072030), the HMS Research Computing Group O2 cluster, and the HMS Research Instrumentation Core Facility. This study was supported by F30 DC017698 (to J.L.), T32 GM007753, and U19 NS104655. R.I.W. and G.M. are HHMI Investigators.

## Online Methods

### Fly husbandry and genotypes

Unless otherwise specified, flies were raised on cornmeal-molasses food (Archon Scientific) in an incubator on a 12-hour:12-hour light:dark cycle at 25°C at 50-70% relative humidity. Flies for the experiments in Figure 2g and Extended Figure 5c were cultured on Nutri-Fly GF German Food (Genessee Scientific) with 0.1% Tegosept (p-hydroxy-benzoic acid, Genessee Scientific), 80 mM propionic acid (Sigma-Aldrich), and 0.6 mM all trans-retinal (ATR; Sigma-Aldrich). Vials containing ATR food were shielded from light with aluminum foil to prevent photoconversion of ATR. The no-ATR control flies for Extended Figure 5c were kept on cornmeal-molasses food. All experiments used flies with at least one wildtype copy of the *white* gene. Genotypes of fly stocks used in each figure are as follows:

<u>Figure 1</u>
*PFNd calcium imaging:*
w/+; P{R16D01-p65.AD}attP40/+; P{R15E01-Gal4.DBD}attP2/PBac{20XUAS-IVS-jGCaMP7f}VK00005
*EPG calcium imaging:*
w/+; +; P{GMR60D05-GAL4}attP2/PBac{20XUAS-IVS-jGCaMP7f}VK00005
*PFNd whole-cell recording:*
w/+; P{R16D01-p65.AD}attP40/ P{20XUAS-IVS-mCD8::GFP}attP40; P{R15E01-Gal4.DBD}attP2/+

<u>Figure 2</u>
*SpsP calcium imaging:*
w/+; P{VT019012-p65.AD}attP40/+; P{R72C10-Gal4.DBD}attP2/ PBac{20XUAS-IVS-jGCaMP7f}VK00005
*LNO2 calcium imaging:*
+; Mi{Trojan-p65AD.2}Vglut[MI04979-Tp65AD.2]/+; P{VT008681-Gal4.DBD}attP2/ PBac{20XUAS-IVS-jGCaMP7f}VK00005
*SpsP optogenetic activation with PFNd whole-cell recording:*
w/+; P{GMR16D01-lexA}attP40/ P{VT019012-p65.AD}attP40; P{13xLexAop2-IVS-pmyr::GFP}VK00005, P{20xUAS-CsChrimson-mCherry-trafficked}su(Hw)attP1/ P{R72C10-Gal4.DBD}attP2

<u>Figure 3:</u>
*PFNv calcium imaging:*
w/+; P{R22G07-p65.AD}attP40/+; P{VT063307-Gal4.DBD}attP2/PBac{20XUAS-IVS-jGCaMP7f}VK00005

<u>Figure 4:</u>
*hΔB calcium imaging:*
+; P{R72B05-p65.AD}attP40/+; P{VT055827-Gal4.DBD}attP2/PBac{20XUAS-IVS-jGCaMP7f}VK00005
*PFNd calcium imaging:*
w/+; P{R16D01-p65.AD}attP40/+; P{R15E01-Gal4.DBD}attP2/PBac{20XUAS-IVS-jGCaMP7f}VK00005

<u>Extended Figure 2:</u>
*PFNd calcium imaging:*
w/+; P{R16D01-p65.AD}attP40/+; P{R15E01-Gal4.DBD}attP2/PBac{20XUAS-IVS-jGCaMP7f}VK00005 *PFNd whole-cell recording:*
w/+; P{R16D01-p65.AD}attP40/ P{20XUAS-IVS-mCD8::GFP}attP40; P{R15E01-Gal4.DBD}attP2/+

<u>Extended Figure 4:</u>
*GFP expression pattern:*
+; Mi{Trojan-p65AD.2}Vglut[MI04979-Tp65AD.2]/P{20XUAS-IVS-mCD8::GFP}attP40; P{VT008681-Gal4.DBD}attP2/+
*MultiColor Flip Out (MCFO):*
+/ w[1118], P{R57C10-FLPL}su(Hw)attP8; Mi{Trojan-p65AD.2}Vglut[MI04979-Tp65AD.2]/+; P{VT008681-Gal4.DBD}attP2/Pbac{10xUAS(FRT.stop)myr::smGdP-HA}VK00005, P{10xUAS(FRT.stop)myr:: smGdP-V5-THS-10xUAS(FRT.stop)myr:: smGdP-FLAG}su(Hw)attP1

<u>Extended Figure 5:</u>
*SpsP optogenetic activation with PFNd whole-cell recording:*
w/+; P{GMR16D01-lexA}attP40/ P{VT019012-p65.AD}attP40; P{13xLexAop2-IVS-pmyr::GFP}VK00005, P{20xUAS-CsChrimson-mCherry-trafficked}su(Hw)attP1/ P{R72C10-Gal4.DBD}attP2
*Empty split-Gal4 optogenetic activation control with PFNd whole-cell recording:*
w/+; P{GMR16D01-lexA}attP40/ P{p65.AD.Uw}attP40; P{13xLexAop2-IVS-pmyr::GFP}VK00005, P{20xUAS-CsChrimson-mCherry-trafficked}su(Hw)attP1/ P{GAL4.DBD.Uw}attP2
*IbSpsP calcium imaging:*
w/+; P{R47G08-p65.AD}attP40/+; P{VT012791-Gal4.DBD}attP2/ PBac{20XUAS-IVS-jGCaMP7f}VK00005

<u>Extended Figure 6:</u>
*PFNv calcium imaging:*
w/+; P{R22G07-p65.AD}attP40/+; P{VT063307-Gal4.DBD}attP2/PBac{20XUAS-IVS-jGCaMP7f}VK00005

<u>Extended Figure 7:</u>
*LNO1 calcium imaging:*
+; P{VT020742-p65.AD}attP40/+; P{VT017270-GAL4.DBD}attP2/ PBac{20XUAS-IVS-jGCaMP7s}VK00005

<u>Extended Figure 8:</u>
*hΔB calcium imaging:*
+; P{R72B05-p65.AD}attP40/+; P{VT055827-Gal4.DBD}attP2/PBac{20XUAS-IVS-jGCaMP7f}VK00005
*PFNd calcium imaging:*
w/+; P{R16D01-p65.AD}attP40/+; P{R15E01-Gal4.DBD}attP2/PBac{20XUAS-IVS-jGCaMP7f}VK00005
*PFNv calcium imaging:*
w/+; P{R22G07-p65.AD}attP40/+; P{VT063307-Gal4.DBD}attP2/PBac{20XUAS-IVS-jGCaMP7f}VK00005

### Origins of transgenic stocks

The following stocks were obtained from the Bloomington Drosophila Stock Center (BDSC) and published as follows: P{GMR60D05-GAL4}attP2 (BDSC 39247)^46^, P{GMR16D01-lexA}attP40 (BDSC 52503)^46^, P{R72B05-p65.AD}attP40 (BDSC 70939)^46^, P{VT055827-Gal4.DBD}attP2 (BDSC 71851)^47^, P{VT008681-Gal4.DBD}attP2 (BDSC 73701)^47^, Mi{Trojan-p65AD.2}Vglut[MI04979-Tp65AD.2] (BDSC 82986)^29^, Pbac{20XUAS-IVS-jGCaMP7f}VK00005 (BDSC 79031)^23^, and P{p65.AD.Uw}attP40; P{GAL4.DBD.Uw}attP2 (BDSC 79603)^48^.

P{20XUAS-IVS-mCD8::GFP}attP40 was a gift from B. Pfeiffer and G. Rubin and was described previously^46^.

The split-Gal4 line targeting PFNd neurons was ss00078 (P{R16D01-p65.AD}attP40; P{R15E01-Gal4.DBD}attP2). The split-Gal4 line targeting SpsP neurons was ss52267 (P{VT019012-p65.AD}attP40; P{R72C10-Gal4.DBD}attP2). The split-Gal4 line targeting IbSpsP neurons was ss04778 (P{R47G08-p65.AD}attP40; P{VT012791-Gal4.DBD}attP2). The split-Gal4 line targeting PFNv neurons was ss52628 (P{R22G07-p65.AD}attP40;P{VT063307-Gal4.DBD}attP2). The split-Gal4 line targeting LNO1 neurons was ss47398 (P{VT020742-p65.AD}attP40; P{VT017270-GAL4.DBD}attP2). These lines were obtained from the Janelia Research Campus FlyBank and have been described previously^3^.

We constructed a split-Gal4 line to target LNO2 neurons that incorporates the Vglut^AD^ transgene^29^. This split-Gal4 line is Mi{Trojan-p65AD.2}Vglut[MI04979-Tp65AD.2]; P{VT008681-Gal4.DBD}attP2. We validated the expression of this line using immunohistochemical anti-GFP staining, and also using Multi-Color-Flip-Out (MCFO) to visualize single-cell morphologies. On occasion, this split line labels a cell type innervating nodulus subunit 3 (NO3); MCFO results suggest that this is a separate cell type from LNO2 and does not innervate NO2 (Extended Data Fig. 4).

The recombinant chromosome P{13xLexAop2-IVS-pmyr::GFP}VK00005, P{20xUAS-CsChrimson-mCherry-trafficked}su(Hw)attP1 was a gift from Vivek Jayaraman’s lab.

MCFO experiments used w[1118], P{R57C10-FLPL}su(Hw)attP8; +; Pbac{10xUAS(FRT.stop)myr::smGdP-HA}VK00005, P{10xUAS(FRT.stop)myr::smGdP-V5-THS-10xUAS(FRT.stop)myr::smGdP-FLAG}su(Hw)attP1 (BDSC 64087)^49^.

We constructed a split-Gal4 line to target hΔB neurons. This split-Gal4 line is +; P{R72B05-p65.AD}attP40; P{VT055827-Gal4.DBD}attP2.

### Fly preparation and dissection

For calcium imaging experiments, we used female flies 20-50 hours post-eclosion and food-deprived (providing only a Kimwipe with water) for at least 12 hours prior to the experiment. No circadian restriction was imposed for the time of experiments. For optogenetic activation experiments in Fig. 2g and Extended Figure 5c, we used female flies 1-5 days post-eclosion. Flies were kept on Nutri-Fly GF German Food with 0.6 mM ATR. For all other electrophysiology experiments, we used female flies 24-48 hours old; 5/7 flies included in our dataset were food-deprived for 12-24 hours. No circadian restriction was imposed for the time of experiments.

Flies were briefly cold anesthetized prior to dissection. For calcium imaging experiments and electrophysiology experiments during walking behavior, we secured the fly in an inverted pyramidal platform CNC-machined from black Delrin (Autotiv, Protolabs) with the head pitched forward so that the posterior surface of the head was more accessible to the microscope objective. For electrophysiology experiments with optogenetic activation, we used a photochemically-etched, flat stainless-steel shim stock platform (Etchit), and the head was oriented normally (dorsal-side up). The wings were removed, and the fly head and thorax were secured to the holder using UV-curable glue (Loctite AA 3972) and cured with ultraviolet light (LED-200, Electro-Lite Co). To remove large brain movements, the proboscis was glued using UV-curable glue. The extracellular saline composition was: 103 mM NaCl, 3 mM KCl, 5 mM TES, 8 mM trehalose, 10 mM glucose, 26 mM NaHCO_3_, 1 mM NaH_2_PO_4_, 1.5 mM CaCl_2_, and 4 mM MgCl_2_ (osmolarity 270-275 mOsm). The saline was bubbled with 95% O2 and 5% CO_2_ to reach a final pH of ~7.3. A window was opened in the head cuticle, and trachea and fat were removed to expose the brain. To further reduce brain movement, muscle 16 was inactivated by gently tugging or clipping the esophagus posteriorly, or by clipping the muscle anteriorly. For electrophysiology experiments, the perineural sheath was removed with fine forceps over the brain region of interest. For all electrophysiology experiments, saline was continuously superfused over the brain; for calcium imaging, saline was superfused prior to experiments.

### Two-photon calcium imaging

We used a galvo-galvo-resonant two-photon microscope (Thorlabs Bergamo II, Vidrio RMR Scanner) with a fast piezoelectric objective scanner (Physik Instrument P725) and a 20×/1.0 NA objective (XLUMPLFLN20XW, Olympus) for volumetric imaging. We used a Chameleon Vision-S Ti-Sapphire femtosecond laser tuned to 940 nm for two-photon GCaMP excitation. Emission was collected on GaAsP PMT detectors (Hamamatsu) through a 525-nm bandpass filter (Thorlabs). We used ScanImage 2018 software^50^ (Vidrio Technologies) to control the microscope, and imaging data were collected in ScanImage using National Instruments PXIe-6341 hardware.

The imaging region for all experiments was 256×128 pixels, with 12 slices in the z-axis for each volume (3-5 μm per slice) resulting in a ~10 Hz volumetric scanning rate. For PFNd and PFNv imaging experiments, we imaged the PB. For SpsP and IbSpsP imaging experiments, we imaged the PB. For LNO2 imaging experiments, we imaged the NO. For hΔB imaging experiments, we imaged the FB.

### Patch-clamp recordings

Thick-wall filamented borosilicate glass (OD 1.5, ID 0.86 mm, Sutter) pipettes with a resistance range of 9-12 MΩ were pulled using a P-97 Sutter puller. Pipettes were filled with an internal solution^51^ consisting of 140 mM KOH, 140 mM aspartic acid, 1 mM KCl, 10 mM HEPES, 1 mM EGTA, 4 mM MgATP, 0.5 mM Na3GTP, and 13 mM biocytin hydrazide, filtered twice through a 0.22-μm PVDF filter. To visualize the cells for recording, we used a FLIR camera (Chameleon3 CM-U3-13Y3C) mounted on an upright compound microscope (Olympus BX51WI) with a 40× water immersion objective (LUMPlanFLN 40XW, Olympus). We used a 100-W Hg arc lamp (Olympus, U-LH100HG) and an eGFP long-pass filter to detect GFP fluorescence. For optogenetics experiments, the brain was illuminated from below using bright field transmitted light through the microscope condenser to identify cell bodies for recording, which was then turned off prior to optogenetic stimulus delivery. For walking experiments, the fly was illuminated from below using a fiber optic coupled LED (M740F2, Thorlabs) coupled to a ferrule-terminated patch cable (200-μM core, 0.22 n.a., Thorlabs) attached to a fiber optic cannula (200-μM core, 0.22 n.a., Thorlabs). The cannula was glued to the ventral side of the holder and positioned approximately 135° from the front of the fly so as to be unobtrusive to the fly’s visual field. Throughout the experiment, saline bubbled with 95% O2 / 5% CO_2_ was superfused over the fly using a gravity pump at a rate of 2 mL/min. Whole cell recordings were performed using an Axopatch 200B amplifier with a CV-203BU headstage (Molecular Devices). Data were low-pass filtered at 5 kHz and acquired on a NiDAQ PCIe-6363 card (National Instruments) at 20 kHz. The liquid junction potential was corrected by subtracting 13 mV from recorded voltages^52^.

### Spherical treadmill and locomotion measurement

For calcium imaging experiments, flies were positioned on a 9-mm ball made from foam (FR-4615, General Plastics). The ball was painted with a black pattern using model paint (Vallejo Black Model Color Paint). The spherical treadmill consisted of this ball floating on air in a concave hemispherical depression on a plenum 3-D printed from clear acrylic (Autotiv). Medical-grade breathing air was flowed through a hole at the bottom of the depression. The ball was illuminated with a round-board 36 infrared LED lamp (SODIAL). Ball movement was tracked using a video camera (CM3-U3-13Y3M-CS, FLIR) fitted with a macro zoom lens (Tamron 23FM08L 8-mm 1:1.4 lens). The camera faced the ball from the right side of the fly at a 90° angle. We removed one panel of the visual panorama to accommodate the camera view of the ball. The camera frame rate was 50 Hz. Machine vision software (FicTrac v2.0) was used to track the position of the ball^53^. We modified FicTrac to output computed ball position parameters in real time through the Redis publish/subscribe messaging paradigm. We wrote custom Python software to read in FicTrac outputs from Redis and to produce analog voltage signals through a Phidget analog output device (Phidget Analog 4-Output 1002_0B). The forward axis ball displacement, yaw axis ball displacement, gain-modified forward ball displacement (not used for experiments in this study), and gain-modified yaw ball displacement were output through the Phidget analog device. For closed-loop experiments, the gain-modified yaw ball displacement voltage signal was used to update the azimuthal position of the visual cues displayed by the visual panorama. All voltage analog signals were digitized and acquired using NiDAQ PCI-6341 (National Instruments) at 4 kHz. The forward axis, lateral axis, and yaw axis ball movements from FicTrac, along with their timestamps, were recorded by the custom Python software and saved to a HDF5 file for each experiment.

For electrophysiology experiments, the following parameters were altered. The ball was illuminated using a 780 nm mounted LED source (M780L3, Thorlabs). The ball’s movement was tracked using GS3-U3-41C6NIR video camera (FLIR) fitted with an InfiniStix 94-mm 0.5× macro zoom lens. One panel 180° behind the fly was removed to accommodate the camera view of the ball and the light source. FicTrac v2.1 was used to track the position of the ball in real time^53^. We recorded the forward, side, and yaw displacement of the ball via a NiDAQ PCIe-6363 card at 20 kHz. Via built-in serial communication support, we used a custom Python script to output FicTrac parameters to a Phidget analog output device (Phidget Analog 4-Output 1002_0B).

### Visual panorama and visual stimuli

To display visual stimuli, we used a circular panorama built from modular square (8×8 pixel) LED panels^24^. The circular arena was 12 panels in circumference and 2 panels tall. For calcium imaging experiments, we removed one panel 90° to the right of the fly; the bottom panel at that azimuth remained to display stimuli. For electrophysiology experiments, we removed one panel 180° behind the fly. In all experiments, the modular panels contained blue LEDs with peak blue (470 nm) emission; blue LEDs were chosen to reduce overlap with the GCaMP emission spectrum. For calcium imaging experiments, four layers of filters were added in front of the LED arena (Rosco, R381) to further reduce overlap in spectra. A final diffuser layer was placed in front of the filters (SXF-0600, Snow White Light Diffuser, Decorative Films). For electrophysiology experiments, only the diffuser layer was used.

The visual stimulus displayed was a bright 2-pixel-wide vertical bar. The bar’s height was the full 2-panel height of the area (except for 75-105° to the right of the fly, when the bar was 1 full panel in height). The azimuth position of the bar was controlled during closed-loop experiments via the voltage signal from the Phidget device, which was used to convert FicTrac outputs to an analog voltage signal. For calcium imaging experiments, a 0.8× yaw gain was used; this meant that for a given yaw displacement of the ball, the visual cue displacement was 0.8× the ball’s yaw displacement. For electrophysiology experiments, a 1× yaw gain was used.

The visual panorama provided no information about body velocity. It therefore seems likely that the velocity signals we observed were due to proprioceptive feedback and/or motor efference copy. However, it is possible that flies could see the pattern on the spherical treadmill moving as they walked, and thus there may also be a contribution from ventral optic flow signals.

### Experimental trial structure during calcium imaging

For calcium imaging experiments, prior to data collection, all flies were allowed to walk for 5 min in darkness and then at least 10 min in closed loop with the visual cue. For calcium imaging experiments, data were collected in two 300-s trials in closed loop with a bright bar; there was a 5-s interval of darkness between trials. For electrophysiology experiments, flies were given at least 10 minutes of walking in closed loop with the visual cue prior to data collection. Each electrophysiology experiment consisted of 3 continuous 200-s closed loop trials with a 1-s inter-trial interval in darkness.

### Optogenetic stimuli and pharmacology

Optogenetic stimuli were delivered using a Hg lamp and an ET-Cy5 long-pass filter (590-650nm, Chroma), with a power of ~10 mW/mm^2^. A shutter (Uniblitz Electronic) was used to control the light pulse duration. Light pulses (10 ms) were delivered at 4-s inter-pulse intervals, in three sessions of 150 pulses each. In the first session, the extracellular saline contained 1 μM TTX (554412, EMD Biosciences). In the second session, 1μM picrotoxin (CAS 124-87-8, Sigma Aldrich) was added. In the third session, picrotoxin was increased to 100 μM. In no-ATR control experiments, the light pulse was 50 ms long.

### Immunohistochemistry

#### General immunochemistry procedures

Brains were dissected from female flies 2-3 days post-eclosion in Drosophila external saline (see above) and fixed in 4% paraformaldehyde (PFA, Electron Microscopy Sciences) in phosphate-buffered saline (Thermo Fisher Scientific) for 15 min. Brains were then washed with PBS before adding blocking solution containing 5% normal goat serum (NGS, Sigma-Aldrich) in PBST (PBS with 0.44% Triton-X, Sigma-Aldrich) for 20 minutes. Brains were then incubated in primary antibody with blocking solution for 24 hrs at room temperature, washed in PBST, and then incubated in secondary antibody with blocking solution for 24 hrs at room temperature. After a final wash in PBST, brains were mounted using Vectashield (Vector Laboratories) for imaging. For MCFO protocols, a tertiary incubation step for 24 hours at room temperature and wash with PBST was performed prior to mounting. Mounted brains were imaged on a Leica SPE confocal microscope using a 40× oil-immersion objective with 1.3 n.a. Image stacks comprised 100-250 z-slices at a depth of 1 μm per slice. Image resolution was 1024×1024 pixels.

#### Visualizing Gal4 expression patterns

The primary antibody solution contained chicken anti-GFP (1:1000, Abcam) and mouse anti-Bruchpilot (1:30, Developmental Studies Hybridoma Bank, nc82). The secondary antibody solution contained Alexa Fluor 488 goat anti-chicken (1:250, Invitrogen) and Alexa Fluor 633 goat anti-mouse (1:250, Invitrogen).

#### MCFO

The primary antibody solution contained mouse anti-Bruchpilot (1:30, Developmental Studies Hybridoma Bank, nc82), rat anti-FLAG (1:200, Novus Biologicals), and rabbit anti-HA (1:300, Cell Signal Technologies). The secondary antibody solution contained Alexa Fluor 488 goat anti-rabbit (1:250, Invitrogen), ATTO 647 goat anti-rat (1:400, Rockland), and Alexa Fluor 405 goat anti-mouse (1:500, Invitrogen). Tertiary antibody solution contained DyLight 550 mouse anti-V5 (1:500, AbD Serotec).

### Data analysis for imaging and electrophysiology experiments

Calcium imaging data analysis was performed on MATLAB 2018a and 2018b; electrophysiology data analysis was performed on MATLAB 2019b. For calcium imaging data analysis, no flies were excluded from the dataset. Analyses for calcium imaging datasets were parallelized on a high-performance computing cluster (O2 High Performance Compute Cluster, HMS Research Computing Group). For electrophysiology analysis, we excluded experiments if the fly did not sample the full 360-degree heading range or if there was large electrical noise. This occurred in 7/16 cells recorded; we included 9 cells across 7 flies in our dataset.

#### Calcium imaging alignment and processing

Rigid motion correction in the x, y, and z axes was performed for each trial using the NoRMCorre algorithm^54^. Each region of interest (ROI) was defined in a single z-plane. For each ROI, a ΔF/F metric was calculated, with the baseline fluorescence (F) defined as the mean of the bottom 5% of fluorescence values within the given trial (300 s in length). For PB imaging, 16 ROIs were defined, one for each of the 16 glomeruli occupied by PFNd dendrites or PFNv dendrites or EPG axons; these ROIs were drawn based on visible anatomical boundaries. For FB imaging, eight ROIs were defined manually over hΔB neurites to correspond to eight columns spanning the horizontal axis of the FB. ROIs were defined to be of roughly equal width and collectively cover the lateral span of the FB without overlap between ROIs. For NO imaging, an ROI was defined for each the left and right NO2, which were anatomically separable.

#### Processing locomotion data in calcium imaging experiments

The displacement of the spherical treadmill was computed by FicTrac in the yaw and forward directions, output from the Phidget device as a voltage signal, and collected by the NI data acquisition device (DAQ). The FicTrac-computed displacements along the yaw, forward, and lateral (side) axes were also saved directly to an HDF5 file. To get the forward and yaw velocity, the voltage signal from the DAQ was first downsampled (using MATLAB downsample function) to the FicTrac output rate (50 Hz), converted to radians, and unwrapped. A second-order Butterworth low-pass filter was applied to the displacement, and velocity was calculated using the MATLAB gradient function. To get the lateral velocity, the FicTrac outputs saved to the HDF5 file needed to be aligned to the DAQ-collected inputs. To do this, the integrated forward displacement was first linearly interpolated to the time points of the DAQ signal (after downsampling to 50 Hz). The interpolated integrated forward displacement was then low-pass filtered using a second-order Butterworth function, and velocity was calculated using the MATLAB gradient function. The forward velocity computed from the HDF5 file was then aligned to the forward velocity computed using voltage signals from the DAQ using the MATLAB finddelay function. The delay calculated between the HDF5 forward velocity signal and the DAQ-input forward velocity signal was found to be consistent across channels, and the aligned HDF5 forward velocity and DAQ-input forward velocity traces were nearly identical. Moreover, applying the interpolation, smoothing, velocity calculation, and delay adjustment procedure to the HDF5 unwrapped heading signal resulted in a yaw velocity trace nearly identical to the one computed using the DAQ-input voltage signal. Thus, we applied the same procedure to the HDF5 integrated lateral displacement to obtain the lateral velocity. Finally, velocity calculated along all three axes were downsampled to the volumetric imaging rate.

For all analyses, we removed the first 3s of every trial to account for the delay in visual stimulus display. We also removed time periods around starting/stopping transitions to account for jGCaMP7 rise and decay kinetics. Specifically, for each trial, a walking transition ‘cutoff for the total speed of the fly (forward speed + lateral speed + yaw speed) was computed by fitting the speed distribution to a bimodal normal mixture model using maximum likelihood estimation and finding the speed at which contribution of the two normal distributions to the mixture PDF was equal. For cases when the mixture model fit was relatively poor and generated a speed cutoff less than 0.1 rad/s, we used a cutoff value of 0.1 rad/s; when the speed cutoff estimate was greater than 0.5 rad/s, we used a cutoff value of 0.5 rad/s. This speed threshold and an additional requirement that walking and stopping epochs be at least 0.5 s in length were used to determine walking transition time points. For PFN, EPG, IbSpsP, and LNO2 imaging, we removed 2·τ_rise_ after stop→walk transitions and 2·τ_decay_ after walk→stop transitions. For SpsP and LNO2 imaging, this correspondence was flipped. Based on published data^23^, we used τ_rise_=75 ms and τ_decay_=520 ms for jGCaMP7f experiments and τ_rise_=70 ms and τ_decay_=1.69 s for jGCaMP7s experiments. The first 200 ms prior to every transition was also removed in our analyses.

In Figs. 1e, 1f, 2b, 2e, 3b, 4f, and Extended Data Figs. 5f and 7b, velocity traces were lightly smoothed using a 300-s moving average filter for display only.

#### Processing locomotion data in electrophysiology experiments

The displacement of the spherical treadmill was computed by FicTrac in the yaw, forward, and lateral directions, output from the Phidget device as a voltage signal, and collected by the NI data acquisition device (DAQ). The voltage signal from the DAQ was first converted into radians and unwrapped. A second-order Butterworth lowpass filter was then applied to the displacement. The displacement traces were then smoothed using the MATLAB smoothdata lowess function. Displacements were downsampled to 25Hz. Velocity was calculated using the MATLAB gradient function and interpolated up to 1000Hz using the MATLAB resample function with a second-order antialiasing filter. For all analyses, the first and last 500 ms of each trial were removed.

#### Ensemble representation of heading direction

To determine the position of the heading bump in the PB (in PFNd, EPG, IbSpsP, and PFNv neurons), we took the spatial Fourier transform of the ΔF/F across the 16 PB glomeruli at every time point. In order to ensure a period of 8 glomeruli in the spatial Fourier transform, we re-arranged the PB glomeruli for each cell type, following published maps^2^. Specifically, for EPG neurons (in which our driver line does not contain neurites in glomeruli L9 and R9), the PB glomeruli were arranged in the following order: L8-L7-L6-L5-L4-L3-L2-L1-R2-R3-R4-R5-R6-R7-R8-R1. For PFNd, PFNv, and IbSpsP neurons (which do not contain neurites in glomeruli L1 and R1), the arrangement was: L9-L8-L7-L6-L5-L4-L3-L2-R9-R2-R3-R4-R5-R6-R7-R8. We defined the bump position as the phase of the Fourier component at a period of eight glomeruli; we used the sign convention in which positive phase change corresponds to rightward movement of the bump in the protocerebral bridge when viewed posteriorly.

To determine the neural heading coding in the FB, we defined each FB column as representing 1/8 of the full 360° azimuthal space. Using the centers of each bin of heading space and the ΔF/F for the given column as weights, we calculated the population vector average across the eight FB ROI columns for each time point. We defined a positive phase change to be a rightward movement of the bump in the FB when viewed posteriorly.

#### Correlation analysis of heading direction

We calculated the circular-circular correlation coefficient between the visual cue position and the position of the heading bump for each fly. We limited the correlation calculation to periods when the bump amplitude (defined as the maximum ΔF/F – minimum ΔF/F across the glomeruli for each half of PB) was > 0.8. For each cell type, we calculated circular-circular correlations for different cue-bump time lags, and we used the time lag where the correlation was maximal. This lag was 0.1 s for EPG imaging, 0.3 s for PFNd imaging, 0.4 s for PFNv imaging, and 0.4 s for IbSpsP imaging.

#### Normalized bump amplitude

For each half of the PB, we defined the bump amplitude as the maximum ΔF/F – minimum ΔF/F across the eight glomeruli. Then, for each fly, we performed min-max normalization of the bump amplitudes, using the mean of the bottom 5% of bump amplitudes as the minimum and the top 5% of bump amplitudes as the maximum. We performed this rescaling of bump amplitudes for each side of the protocerebral bridge separately. This rescaling enabled us to compare between the right and left halves of the protocerebral bridge and average data across flies.

For the FB, we defined the bump amplitude as the maximum ΔF/F – minimum ΔF/F across the eight columns. Then, for each fly, we performed min-max normalization of the bump amplitudes, using the mean of the bottom 5% of bump amplitudes as the minimum and the top 5% of bump amplitudes as the maximum.

#### Computing population activity in the PB as a function of translational velocity

We binned each time point based on the forward and lateral velocity of the fly. Lateral velocity was defined in the ipsilateral direction (thus right was positive in analyzing the right PB, while left was positive in analyzing the left PB). We then pooled data across the left and right PB. We required that each 2D velocity bin contain at least 10 datapoints (or approximately 1s of data) for a given fly to qualify for inclusion in the group analysis. We then calculated the mean normalized bump amplitude within each velocity bin for every fly. We then took the mean across flies to produce the family of curves. If over half of the flies in the dataset did not have enough observations for a given forward/side velocity bin, we excluded the bin entirely from our dataset; otherwise, we took the mean across the flies that had enough observations to be included in the dataset. In our dataset, bin exclusion only occurred at the ends of the specified bin edges. For these analyses, we used a time lag that produced the maximally steep relationship between activity and velocity. For PFNd and EPG neurons, this time lag was 0.2 s. For PFNv neurons, this time lag was 0.3 s. For IbSpsP neurons, this lag was 0.1 s.

#### Computing population activity in the PB as a function of translational velocity angle

We binned each time point based on the translation angle and translation speed of the fly, where translational angle was calculated as the vector angle and the translation speed as the vector magnitude defined by the vector sum of forward and lateral velocity. A vector angle of zero was defined as aligned with the heading of the fly (i.e., lateral velocity was zero), and positive angles were defined to be to the ipsilateral direction of the population (e.g. for PFNd.R neurons, positive angle was to the right; for PFNd.L, a positive angle was to the left). We then pooled data across the left and right PB. We required that each 2D velocity bin contain at least 10 datapoints (or approximately 1s of data) for a given fly to qualify for inclusion in the group analysis. We then calculated the mean normalized bump amplitude within each velocity bin for every fly. We then took the mean across flies to produce the family of curves. If over half of the flies in the dataset did not have enough observations for a given angle/magnitude bin, we excluded the bin entirely from our dataset; otherwise, we took the mean across the flies that had enough observations to be included in the dataset. Bin exclusion only occurred at the ends of the specified bin edges. We used the same time lags as above.

#### Computing direction of preferred translational velocity

We first calculated the mean normalized PB bump amplitude over binned forward or lateral velocities for each fly (combining the data from the left and right PB by representing lateral velocity in the ipsilateral direction). We then took the mean across flies. For both PFNd and PFNv neurons, these relationships were fairly linear (Extended Data Figs. 2b, 6b). We computed the slope of the linear regression between normalized bump amplitude and forward velocity, as well as the slope of the linear regression between normalized bump amplitude and ipsilateral side velocity (Extended Data Figs. 2b, 6b). To calculate the preferred translational velocity angle, we took the arctangent of the ratio between the slope of the normalized bump amplitude versus lateral velocity and the slope of the normalized bump amplitude versus forward velocity (Extended Data Figs. 2c, 6c). We assumed that the preferred directions of the left and right PB were mirror-symmetric.

#### PFNd electrophysiology analysis during walking behavior

Voltage traces were downsampled to 1000 Hz using the MATLAB resample function. To remove spikes, voltage traces were then median filtered using the medfilt1 MATLAB function. To calculate firing rate, the median filtered trace was subtracted from the downsampled trace to isolate spike times. Timepoints of spikes were identified using the MATLAB findpeaks function on the baselined trace with the minimum peak height specified for every experiment. To estimate firing rate, identified spikes were smoothed using a 10 ms Gaussian kernel. Finally, the voltage trace was further smoothed using the MATLAB smoothdata function to remove remaining noise.

#### Computing PFNd membrane voltage and firing rate as a function of translational velocity

The preferred heading direction of each cell was estimated by generating a heatmap of membrane potential values as a function of heading and forward velocity, in which heading was binned in 10° segments from −180° to 180°. In addition, we generated a second plot of the mean membrane potential value as a function of heading only. We used a preferred translational angle magnitude of 31° for our analyses. To determine the direction of the preferred translational angle (i.e. −31° or + 31°), we generated heatmaps of the membrane potential and firing rate binned by forward velocity and lateral velocity. Once the sign of the preferred translational direction vector offset was determined, velocity in the preferred translational direction was found by projecting the forward and lateral velocities onto this preferred travel direction vector and summing the two components.

To determine the temporal relationship between activity and behavior (Fig. 1j), we used continuous trial segments of at least 30 seconds during which: (1) at least 60% of the fly’s heading fell within a 120° range around the preferred heading, and (2) there were multiple blocks of non-zero velocity. We used the MATLAB xcorr function to determine the cross correlation between changes in the membrane potential of the cell and changes in the fly’s velocity in the preferred translational direction. A peak in the cross correlation at negative time values indicates that changes in the membrane potential precede changes in velocity; a peak at positive values indicates that changes in the membrane potential follow changes in velocity.

To determine the relationship between activity and velocity in the preferred translation direction (Fig. 1k), we computed membrane voltage and firing rate as a function of velocity along preferred translational direction, binning the data by heading to account for the heading tuning of the cell. We used heading bins of +/− 60° to the preferred heading of the cell and +/− 60° opposite to the preferred heading of the cell. We shifted the behavioral data forward by 113s to account for the behavior-neuron lag; this value was chosen because it was the median lag of the cross-correlation peak across all cells (as shown in Fig. 1j).

#### Computing SpsP, LNO2, and LNO1 activity as a function of translational velocity

We combined data for the left and right hemisphere and binned each time point based on forward and lateral velocity (in the ipsilateral direction). For each fly, we required that a given 2D velocity bin contain at least 10 datapoints (or approximately 1s of data) for inclusion. We then calculated the mean ΔF/F within each velocity bin. To produce the family of curves in Figs. 2c, 2f, and Extended Data Figs. 7c, we took the mean across flies for each velocity bin. A velocity bin was only included in the figure if every fly in the dataset had the minimum required number of datapoints for that bin. Bin exclusion only occurred at the ends of the specified bin edges. For these analyses, we used a time lag that produced the maximally steep relationship between activity and velocity. For SpsP the lag was 0.2 s; for LNO2, the lag was 0.1 s; for LNO1, the lag was 0.2 s.

#### Optogenetic stimulation during patch-clamp recording

In Fig. 2g, the response to the optogenetic stimulus in each pharmacological condition was averaged over 50-90 trials where the response was stable. The pre-stimulus baseline was defined as the mean voltage in the window 2-s prior to the stimulus. The stimulus response was taken as the maximum voltage deviation from baseline within 200 ms after stimulus onset. We found that differences in inhibition are significant comparing TTX condition to TTX + 100 μM picrotoxin (P=2.67×10^−4^, paired-sample t-test with Bonferroni-corrected α = 0.0167, Bonferroni-corrected CI = [−7.44, −3.28] mV), and comparing to TTX + 1 μM picrotoxin condition to TTX + 100 μM picrotoxin condition (P=7.02×10^−4^, paired-sample t-test with Bonferroni-corrected α = 0.0167, Bonferroni-corrected CI = [−7.03, −2.49] mV). Differences in inhibition are not significant comparing TTX condition with TTX + 1 μM picrotoxin condition (P=.1906, paired-sample t-test with Bonferroni-corrected α = 0.0167, Bonferroni-corrected CI = [−2.02, 0.81] mV). Statistical testing used all 6 experiments where each pharmacological treatment was tested.

#### Computing hΔB population activity as a function of translational velocity

We binned each timepoint by the translational speed of the fly, defined as the magnitude of the vector sum of the forward and lateral velocity. We then calculated the mean normalized FB bump amplitude within each speed bin for every fly. We also calculated the mean across flies. We required that all flies contain at least 10 datapoints within a given speed bin for inclusion; this only occurred at the bin edges. The same procedure was used to compute the relationship between normalized bump amplitude and forward velocity and lateral speed. For hΔB neurons, we used a lag of 0.2 s.

#### Computing bump position shift from heading as a function of translational angle

For PFNd and PFNv neurons, we defined the bump position as the phase of the spatial Fourier transform at a period of eight glomeruli (see above). For hΔB neurons, we defined the bump position as the population vector average across the eight columns (see above). We computed a difference value defined as (cue position – bump position), mean centered across the experiment. Mean centering was performed because the heading representation in the EB and PB is arbitrary relative to the cue position^6^. In heading-only representations, the bump position should move to the right the same amount as the cue moves to the right, which occurs when the fly is rotating to the left. Positive difference values indicate that the bump position is further to the left than the cue position, and negative difference values indicate that the bump position is further to the right than the cue position. We also computed the translation angle as the arctangent of the lateral velocity and the forward velocity; positive angles denote rightward translation. For all cell types, we used a lag value of 0.2 s. For each fly, we displayed a histogram of offset values binned by translation angle. We omitted the 90° translation angle bin representing backwards walking because that bin was sparsely sampled. For each translation angle bin, we found the circular mean for each histogram, which is plotted in Fig. 4j and 4k. In Fig. 4j (hΔB neurons), we found the shift was significant when comparing left translation-heading deviations to centered translation-heading deviations (P=0.0030, paired-sample t-test with Bonferroni-corrected α = 0.0167, Bonferroni-corrected CI = [−0.478, −0.140] radians) and when comparing right translation-heading deviations to centered translation-heading deviations (P=0.0038, paired-sample t-test with Bonferroni-corrected α = 0.0167, Bonferroni-corrected CI = [−0.411, −0.105] radians). In Fig. 4k (PFNd neurons), the shift is not significant when comparing left translation-heading deviations to centered translation-heading deviations (P=0.0909, paired-sample t-test with Bonferroni-corrected α = 0.0167, Bonferroni-corrected CI = [−0.1329, 0.0262] radians) or when comparing right translation-heading deviations to centered translation-heading deviations (P=0.0446, paired-sample t-test with Bonferroni-corrected α = 0.0167, Bonferroni-corrected CI = [−0.0156, 0.1520] radians). In Extended Data Fig. 8e (PFNv neurons), this shift is significant when comparing left translation-heading deviations to centered translation-heading deviations (P=0.0084, paired-sample t-test with Bonferroni-corrected α = 0.0167, Bonferroni-corrected CI = [0.0148, 0.2107] radians) and when comparing right translation-heading deviations to centered translation-heading deviations (P=0.0030, paired-sample t-test with Bonferroni-corrected α = 0.0167, Bonferroni-corrected CI = [0.0332,0.204] radians) – note that the shift is in the opposite direction than that for hΔB neurons.

### Connectomics analysis

Using the partial connectome of the adult female fly brain (hemibrain v1.1)^9^, we first obtained the neuron IDs for all hΔB neurons in the dataset (19 in total), and we performed a neuprint^55^ Common Input search to obtain all the inputs to all hΔB neurons. We then discarded all neurons not identified as PFNd or PFNv to obtain a connectivity matrix, which tabulated the number of synapses connecting all PFNd/PFNv neurons to all hΔB neurons.

For Fig. 3h, we assigned each hΔB axon to one of 12 vertical FB columns. We then assigned each PFNd and PFNv neuron to one of 8 vertical FB columns. Using the column designations, we determined the relative lateral location of each neuron along the FB. Using the connectivity matrix, we evaluated every synaptic connection between an hΔB neuron and a PFNd or PFNv neuron by calculating the relative distance between the hΔB axon and the PFNd or PFNv axon. If the distance between the PFN axon and the hΔB axon was <20% of FB’s horizontal extent, we categorized the synapses between the pair as axo-axonal. If the distance between PFN axon and hΔB axon was 30-70% of the FB’s horizontal extent, we categorized the synapses as axodendritic. Only two synapses from this analysis were indeterminate, and we manually classified as axodendritic.

For Fig. 3j, we assigned each PFNd and PFNv neuron a relative preferred heading, based on its cognate PB glomerulus. For example, neurons in L5 or R5 were assigned a preferred heading of 0°, while neurons in L7 and R3 were assigned a preferred heading of +90°. We used the connectivity matrix to determine the input weight of each PFNd and PFNv neuron onto each hΔB neuron. We then computed the connectivity-weighted circular mean of these preferred headings, for each hΔB neuron and each input population (PFNd.right, PFNd.left, PFNv.right, PFNv.left).

For Extended Data Fig. 3, we performed a neuprint Common Input search to obtain all the inputs to PFNd and PFNv neurons. We grouped input synapses by cell type and discarded synapses where the input cell type was undefined or where the cell type made three or fewer synapses across all PFNd/PFNv neurons. We then plotted the distribution of input synapses across cell type using the Total Weight values. In addition, for PFNd neurons, we created a connectivity matrix between PFNd neurons and all input neurons belonging to the top ten input cell types. We discarded any input weight values of three or fewer.

For Extended Data Fig. 4c, we used neuprintr and natverse software^56^ to display the LNO2 skeleton from the hemibrain dataset.

### Computational model

The model was built based on experimentally estimated number, connectivity, and activity of three populations of neurons: PFNd, PFNv, and hΔB. The model comprised 40 PFNd neurons, 20 PFNv neurons, and 19 hΔB neurons whose activity is given by the following equations:

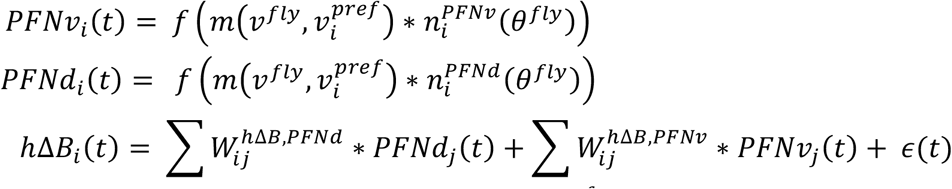

where *v^fly^* is the current body-centric velocity vector and *v^pref^* and the preferred direction of the population. *f* denotes a monotonic non-linearity. For simplicity, we take *f* to be a threshold-linear function. The form of *m* was chosen to approximate measured activity and for simplicity kept the same across both PFN populations (but with differing preferred velocity direction). It is given by:

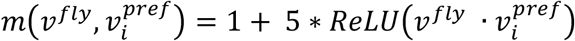

Where · corresponds to a dot product and ReLU is a function that equals *x* if *x*≥0 and 0 otherwise. The dependence of PFN activity on current heading *n* approximates the inheritance of heading tuning through the protocerebral bridge and is given by:

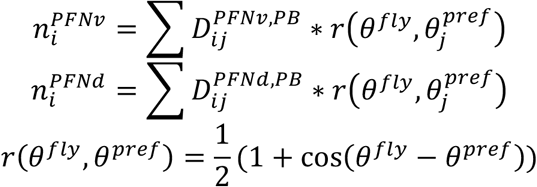

where *r* corresponds to a simplified description of heading tuning in the protocerebral bridge, with *θ^fly^* the fly’s current heading. The matrices *D* are based on anatomical data and map PFN neurons to the protocerebral bridge. The structure of connections from the PFNd and PFNv populations to hΔB in the model were taken directly from the hemibrain connectome^9^, based on the assumption that functional connection weights scale with the number of synapses per connection^57^; these weights were then scaled uniformly by a single, positive scalar value to generate the connectivity matrices *W. ϵ* denotes Gaussian output noise. Simulations were performed in Python.

### Data availability

The datasets generated during and/or analyzed during the current study are available from the corresponding author on reasonable request.

### Code availability

Code for implementing the computational model will be posted to GitHub upon publication.

## References

1. Bicanski, A. & Burgess, N. Neuronal vector coding in spatial cognition. Nat Rev Neurosci 21, 453–470 (2020).

2. Wolff, T., Iyer, N. A. & Rubin, G. M. Neuroarchitecture and neuroanatomy of the *Drosophila* central complex: A GAL4-based dissection of protocerebral bridge neurons and circuits. J. Comp. Neurol. 523, 997–1037 (2015).

3. Wolff, T. & Rubin, G. M. Neuroarchitecture of the Drosophila central complex: A catalog of nodulus and asymmetrical body neurons and a revision of the protocerebral bridge catalog. J Comp Neurol 526, 2585–2611 (2018).

4. Namiki, S. & Kanzaki, R. Comparative neuroanatomy of the lateral accessory lobe in the insect brain. Front. Physiol. 7 (2016).

5. Namiki, S., Dickinson, M. H., Wong, A. M., Korff, W. & Card, G. M. The functional organization of descending sensory-motor pathways in *Drosophila*. eLife 7, e34272 (2018).

6. Seelig, J. D. & Jayaraman, V. Neural dynamics for landmark orientation and angular path integration. Nature 521, 186–191 (2015).

7. Green, J., Adachi, A., Shah, K. K., Hirokawa, J. D., Magani, P. S. & Maimon, G. A neural circuit architecture for angular integration in *Drosophila*. Nature 546, 101–106 (2017).

8. Turner-Evans, D., Wegener, S., Rouault, H., Franconville, R., Wolff, T., Seelig, J. D., Druckmann, S. & Jayaraman, V. Angular velocity integration in a fly heading circuit. eLife 6 (2017).

9. Scheffer, L. K., Xu, C. S., Januszewski, M., Lu, Z., Takemura, S. Y., Hayworth, K. J., Huang, G. B., Shinomiya, K., Maitlin-Shepard, J., Berg, S. et al. A connectome and analysis of the adult Drosophila central brain. eLife 9 (2020).

10. Kim, I. S. & Dickinson, M. H. Idiothetic path integration in the fruit fly *Drosophila melanogaster*. Curr. Biol. 27, 2227–2238 e2223 (2017).

11. Müller, M. & Wehner, R. Path integration in desert ants, Cataglyphis fortis. Proc Natl Acad Sci U S A 85, 5287–5290 (1988).

12. Esch, H. & Burns, J. Distance estimation by foraging honeybees. J Exp Biol 199, 155–162 (1996).

13. DeAngelis, B. D., Zavatone-Veth, J. A. & Clark, D. A. The manifold structure of limb coordination in walking *Drosophila*. eLife 8 (2019).

14. Geurten, B. R., Jahde, P., Corthals, K. & Gopfert, M. C. Saccadic body turns in walking Drosophila. Front Behav Neurosci 8, 365 (2014).

15. Wittlinger, M., Wehner, R. & Wolf, H. The ant odometer: stepping on stilts and stumps. Science 312, 1965–1967 (2006).

16. Ronacher, B. D. & Wehner, R. Desert ants *Cataglyphis fortis* use self-induced optic flow to measure distance travelled. Journal of Comparative Physiology A 177, 21–27 (1995).

17. Schöne, H. Optokinetic speed control and estimation of travel distance in walking honeybees. Journal of Comparative Physiology A 179, 587–592 (1996).

18. Heinze, S., Narendra, A. & Cheung, A. Principles of insect path integration. Curr. Biol. 28, R1043–R1058 (2018).

19. Hulse, B. K., Haberkern, H., Franconville, R., Turner-Evans, D. B., Takemura, S., Wolff, T., Noorman, M., Dreher, M., Dan, C., Parekh, R., Hermundstad, A. M., Rubin, G. M. & Jayaraman, V. A connectome of the Drosophila central complex reveals network motifs suitable for flexible navigation and context-dependent action selection. bioRxiv, doi:10.1101/2020.1112.1108.413955 (2020).

20. Okubo, T. S., Patella, P., D’Alessandro, I. & Wilson, R. I. A neural network for wind-guided compass navigation. Neuron 107, 924–940 e918 (2020).

21. Hardcastle, B. J., Omoto, J. J., Kandimalla, P., Nguyen, B.-C. M., Keleş, M. J., N.K., B., Hartenstein, V. & Frye, M. A. A visual pathway for skylight polarization processing in Drosophila. bioRxiv, doi.org/10.1101/2020.1109.1110.291955 (2020).

22. Stone, T., Webb, B., Adden, A., Weddig, N. B., Honkanen, A., Templin, R., Wcislo, W., Scimeca, L., Warrant, E. & Heinze, S. An anatomically constrained model for path integration in the bee brain. Curr. Biol. 27, 3069–3085.e3011 (2017).

23. Dana, H., Sun, Y., Mohar, B., Hulse, B. K., Kerlin, A. M., Hasseman, J. P., Tsegaye, G., Tsang, A., Wong, A., Patel, R. et al. High-performance calcium sensors for imaging activity in neuronal populations and microcompartments. Nat Methods 16, 649–657 (2019).

24. Reiser, M. B. & Dickinson, M. H. A modular display system for insect behavioral neuroscience. J. Neurosci. Methods 167, 127–139 (2008).

25. Rayshubskiy, A., Holtz, S. L., D’Alessandro, I., Li, A. A., Vanderbeck, Q. X., Haber, I. S., Gibb, P. W. & Wilson, R. I. Neural circuit mechanisms for steering control in walking Drosophila. bioRxiv 2020.04.04.024703; doi.org/10.1101/2020.04.04.024703 (2020).

26. Zheng, Z., Lauritzen, J. S., Perlman, E., Robinson, C. G., Nichols, M., Milkie, D., Torrens, O., Price, J., Fisher, C. B., Sharifi, N. et al. A complete electron microscopy volume of the brain of adult *Drosophila* melanogaster. Cell 174, 730–743 (2018).

27. Eckstein, N., Bates, A. S., Du, M., Hartenstein, V., Jefferis, G. S. X. E. & Funke, J. Neurotransmitter classification from electron microscopy images at synaptic sites in Drosophila. bioRxiv, doi.org/10.1101/2020.1106.1112.148775 (2020).

28. Liu, W. W. & Wilson, R. I. Glutamate is an inhibitory neurotransmitter in the Drosophila olfactory system. Proc Natl Acad Sci U S A 110, 10294–10299 (2013).

29. Lacin, H., Chen, H. M., Long, X., Singer, R. H., Lee, T. & Truman, J. W. Neurotransmitter identity is acquired in a lineage-restricted manner in the Drosophila CNS. eLife 8 (2019).

30. Eschbach, C., Fushiki, A., Winding, M., Schneider-Mizell, C. M., Shao, M., Arruda, R., Eichler, K., Valdes-Aleman, J., Ohyama, T., Thum, A. S. et al. Recurrent architecture for adaptive regulation of learning in the insect brain. Nat Neurosci 23, 544–555 (2020).

31. Wanner, A. A. & Friedrich, R. W. Whitening of odor representations by the wiring diagram of the olfactory bulb. Nat Neurosci 23, 433–442 (2020).

32. Wittman, T. & Schwegler, H. Biol. Cybern. 73, 569–575 (1995).

33. Homberg, U., Heinze, S., Pfeiffer, K., Kinoshita, M. & el Jundi, B. Central neural coding of sky polarization in insects. Philos Trans R Soc Lond B Biol Sci 366, 680–687 (2011).

34. Srinivasan, M., Zhang, S., Lehrer, M. & Collett, T. Honeybee navigation en route to the goal: visual flight control and odometry. J Exp Biol 199, 237–244 (1996).

35. Vinepinsky, E., Cohen, L., Perchik, S., Ben-Shahar, O., Donchin, O. & Segev, R. Representation of edges, head direction, and swimming kinematics in the brain of freely-navigating fish. Sci. Rep. 10, 14762 (2020).

36. Wang, C., Chen, X., Lee, H., Deshmukh, S. S., Yoganarasimha, D., Savelli, F. & Knierim, J. J. Egocentric coding of external items in the lateral entorhinal cortex. Science 362, 945–949 (2018).

37. Hinman, J. R., Chapman, G. W. & Hasselmo, M. E. Neuronal representation of environmental boundaries in egocentric coordinates. Nat Commun 10, 2772 (2019).

38. Alexander, A. S., Carstensen, L. C., Hinman, J. R., Raudies, F., Chapman, G. W. & Hasselmo, M. E. Egocentric boundary vector tuning of the retrosplenial cortex. Sci Adv 6, eaaz2322 (2020).

39. Solstad, T., Boccara, C. N., Kropff, E., Moser, M. B. & Moser, E. I. Representation of geometric borders in the entorhinal cortex. Science 322, 1865–1868 (2008).

40. Savelli, F., Yoganarasimha, D. & Knierim, J. J. Influence of boundary removal on the spatial representations of the medial entorhinal cortex. Hippocampus 18, 1270–1282 (2008).

41. Deshmukh, S. S. & Knierim, J. J. Influence of local objects on hippocampal representations: Landmark vectors and memory. Hippocampus 23, 253–267 (2013).

42. Lever, C., Burton, S., Jeewajee, A., O’Keefe, J. & Burgess, N. Boundary vector cells in the subiculum of the hippocampal formation. J Neurosci 29, 9771–9777 (2009).

43. Byrne, P., Becker, S. & Burgess, N. Remembering the past and imagining the future: a neural model of spatial memory and imagery. Psychol Rev 114, 340–375 (2007).

44. Bicanski, A. & Burgess, N. A neural-level model of spatial memory and imagery. eLife 7 (2018).

45. Lyu, C., Abbott, L. F. & Maimon, G. A neuronal circuit for vector computation builds an allocentric traveling-direction signal in the Drosophila fan-shaped body. bioRxiv, see companion manuscript (2020).

## Additional references for Online Methods

46. Pfeiffer, B., Ngo, T.-T. B., Hibbard, K. L., Murphy, C., Jenett, A., Truman, J. W. & Rubin, G. M. Refinement of tools for targeted gene expression in *Drosophila*. Genetics 186, 735–755 (2010).

47. Tirian, L. & Dickson, B. J. The VT GAL4, LexA, and split-GAL4 driver line collections for targeted expression in the *Drosophila* nervous system. bioRxiv, 198648 (2017).

48. Dionne, H., Hibbard, K. L., Cavallaro, A., Kao, J. C. & Rubin, G. M. Genetic reagents for making split-GAL4 lines in Drosophila. Genetics 209, 31–35 (2018).

49. Nern, A., Pfeiffer, B. D. & Rubin, G. M. Optimized tools for multicolor stochastic labeling reveal diverse stereotyped cell arrangements in the fly visual system. Proc. Natl. Acad. Sci. U. S. A. 112, E2967–2976 (2015).

50. Pologruto, T. A., Sabatini, B. L. & Svoboda, K. ScanImage: flexible software for operating laser scanning microscopes. Biomed. Eng. Online 2, 13 (2003).

51. Wilson, R. I. & Laurent, G. Role of GABAergic inhibition in shaping odor-evoked spatiotemporal patterns in the Drosophila antennal lobe. J. Neurosci. 25, 9069–9079 (2005).

52. Gouwens, N. W. & Wilson, R. I. Signal propagation in Drosophila central neurons. J. Neurosci. 29, 6239–6249 (2009).

53. Moore, R. J., Taylor, G. J., Paulk, A. C., Pearson, T., van Swinderen, B. & Srinivasan, M. V. FicTrac: a visual method for tracking spherical motion and generating fictive animal paths. J. Neurosci. Methods 225, 106–119 (2014).

54. Pnevmatikakis, E. A. & Giovannucci, A. NoRMCorre: An online algorithm for piecewise rigid motion correction of calcium imaging data. J. Neurosci. Methods 291, 83–94 (2017).

55. Clements, J., Dolafi, T., Umayam, L., Neubarth, N. L., Berg, S., Scheffer, L. K. & Plaza, S. M. neuPrint: analysis tools for connectomics. bioRxiv, doi.org/10.1101/2020.1101.1116.909465 (2020).

56. Bates, A. S., Manton, J. D., Jagannathan, S. R., Costa, M., Schlegel, P., Rohlfing, T. & Jefferis, G. S. X. E. The natverse, a versatile toolbox for combining and analysing neuroanatomical data. eLife 9, e53350 (2020).

57. Tobin, W. F., Wilson, R. I. & Lee, W. A. Wiring variations that enable and constrain neural computation in a sensory microcircuit. eLife 6 (2017).

