## Supplemental Figures for "Transforming representations of movement from body- to world-centric space"

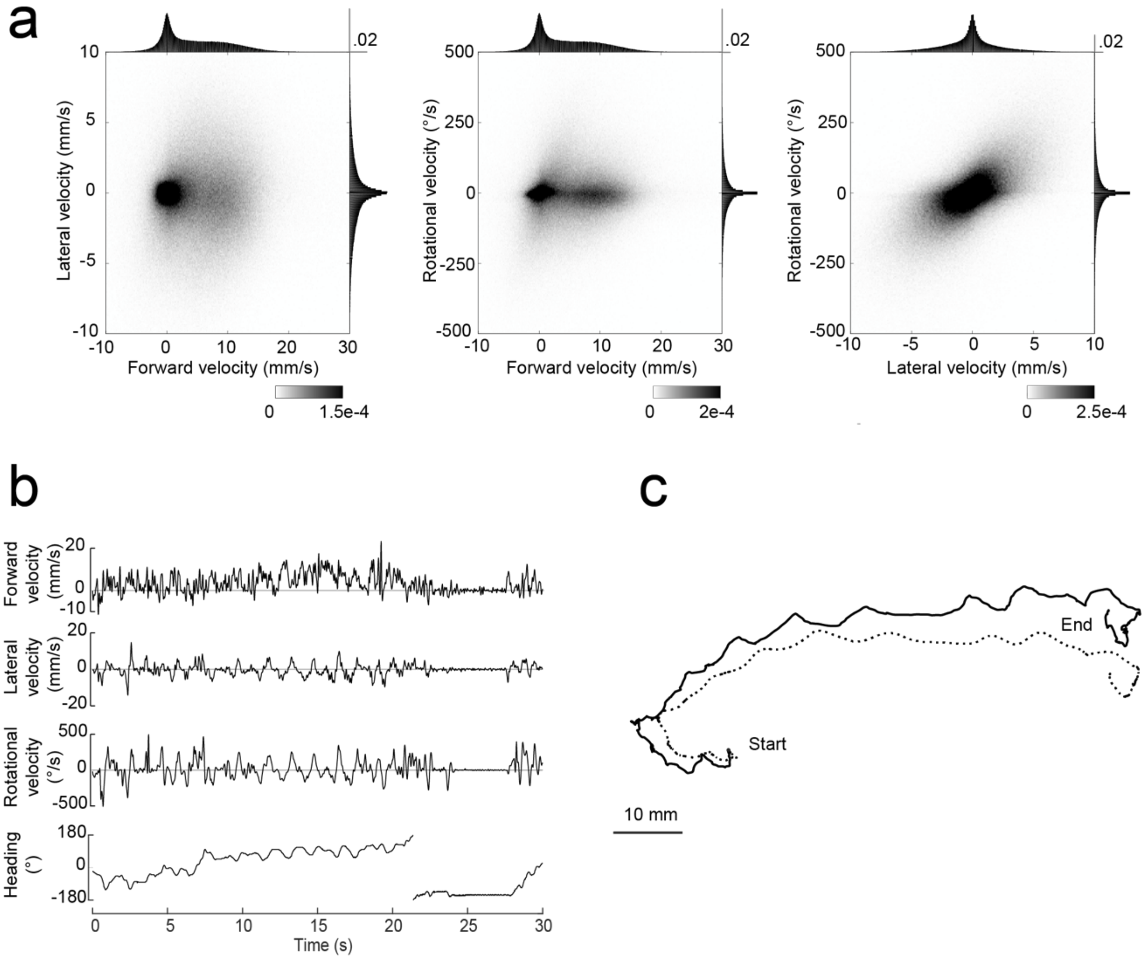


**Extended Data Figure 1: Walking statistics on a spherical treadmill.**

a. Distribution of the forward × lateral, forward × rotational, and lateral × rotational velocities. Shown along each axis is the marginal distribution (gray lines on top right of each heatmap denote scale for the marginal distribution). Data is pooled across n=27 flies. We used the velocities recorded at the camera sampling rate (50 Hz) prior to down-sampling to volumetric calcium imaging rate.

b. An example walking bout (30 seconds). Shown are the fly's forward, lateral, and rotational velocity as well as its heading (based on the position of the visual cue shown in closed loop; note that we used a visual closed loop gain of 0.8×, meaning that the landmark is displaced by an azimuthal angle equal to 0.8× the ball's yaw displacement).

c. Fictive trajectory of the fly in two-dimensional space based on the walking parameters in the example bout shown in b. The dotted line shows the calculated trajectory using only the forward velocity and the heading of the fly, ignoring the lateral velocity. The solid line shows the calculated trajectory using the forward velocity, lateral velocity, and heading of the fly. Note that the dotted line underestimates the curvature of the fly's path.

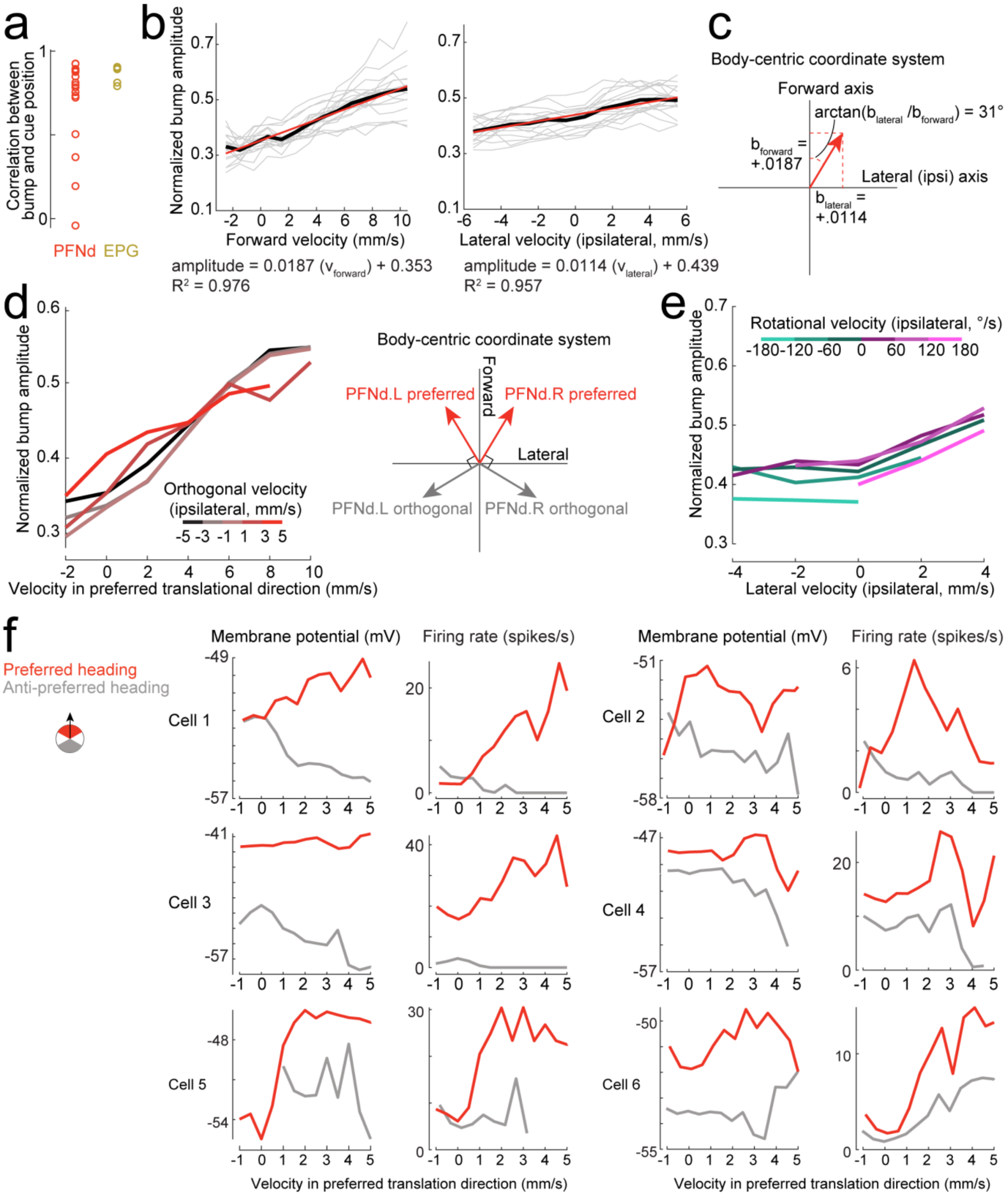


**Extended Data Figure 2: PFNd tuning properties.**

a. Circular correlation between bump and cue position for PFNd (n=16 flies) and EPG neurons (n=5 flies).

b. Normalized PFNd PB bump amplitude versus forward velocity (left), and lateral velocity (right). Gray lines are individual flies and the black line is the mean across flies (n=16 flies). Data from the right and left PB are combined, and lateral velocity is computed in the ipsilateral direction. The red line shows the linear fit to the mean line, with the fitted equation below each plot.

c. Computation of preferred translational direction angle using the linear regression slopes for forward and lateral velocity. We used ratio of the slopes of the linear fits to lateral and forward velocity to calculate the angle of preferred translational direction.

d. Normalized PFNd bump amplitude versus velocity in the preferred translational direction (left). Data from the right and left PB are combined and binned by the fly's velocity orthogonal to the preferred translational direction. Shown is the mean across flies (n=16 flies). Note that a positive value in the orthogonal axis is in the ipsilateral direction.

e. Normalized PFNd bump amplitude versus lateral velocity in the ipsilateral direction. Data from the right and left PB are combined, binned by ipsilateral rotational velocity, and averaged across flies (n=16 flies). Because rotational and lateral velocity are correlated, rotational velocity bins are asymmetrically populated. This analysis shows that there is little or no systematic relationship between PFNd activity and rotational velocity, once we account for the effect of lateral velocity.

f. Same as Fig. 1k, but for several example PFNd cells. We consistently observed that increasing velocity (in the preferred direction) caused depolarization around the neuron's preferred heading, but hyperpolarization around the neuron's anti-preferred heading. Cell 1 is the same cell shown in Fig. 1k, reproduced here for comparison.

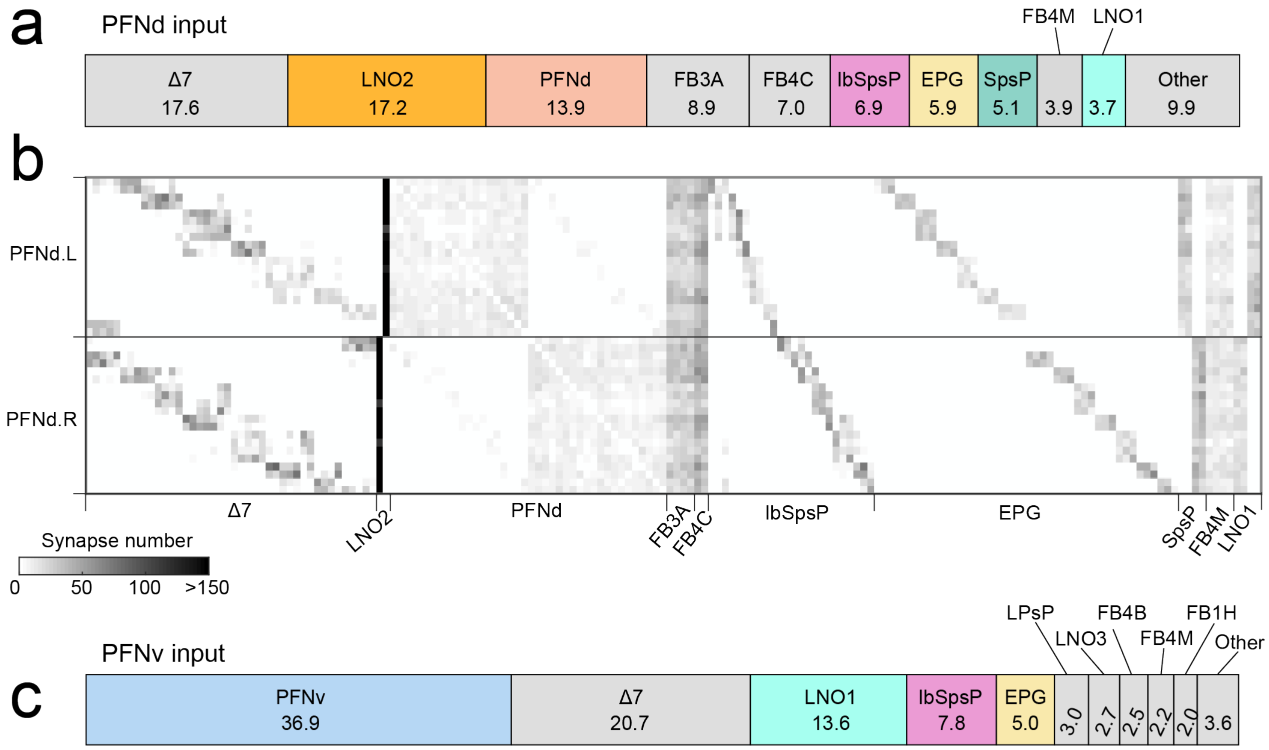


**Extended Data Figure 3: Connectomics analysis of inputs to PFNd and PFNv neurons.**

a. Distribution of input synapses onto PFNd neurons from the hemibrain connectome[9], grouped by cell type. Shown are the top ten cell type inputs onto PFNd neurons; all other identified cell types are grouped into "Other." Collectively, the distribution shown comprises 94.2% of all input synapses onto PFNd neurons. Numbers indicate the percentage of synapses contributed by each input cell type. Note that Δ7 neurons and FB3A/4C/4M neurons are major inputs to PFNd neurons, but we did not screen these neurons as part of our search for the origin of body-centric velocity signals in PFNd neurons, for the following reasons:

Δ7 neurons: Δ7 population activity is known to encode the fly's heading direction, reflecting the strong input to Δ7 neurons from EPG neurons. It has been proposed that the function of Δ7 neurons is to reshape the heading bump into a cosine-shaped activity profile[19]. Thus, much of the "compass input" that we refer to in our study as originating from EPG neurons is probably due to the combined action of EPG neurons (which constitute the primary computational map of the compass system) and Δ7 neurons (which reshape and reinforce the compass system output).

FB3A/4C/4M neurons: These neurons are FB tangential cells, meaning their axons run across the entire horizontal extent of the FB, perpendicular to PFNd dendrites[19]. Like other FB tangential cells, these neurons receive input from outside the central complex and they synapse onto a variety of cell types in the FB. There is evidence that FB tangential cells encode information about context, behavioral state, and internal physiological needs, including the need for sleep[19].

b. Input connectivity matrix for PFNd neurons, shown for the top ten input cell types. Connections comprising 3 or fewer synapses are not shown. Note that the cell types that provide major unilateral input to PFNd neurons are LNO2, IbSpsP, EPG, SpsP, and LNO1.

c. Same as (a) but for PFNv neurons. Collectively, the distribution shown comprises 93.1% of all input synapses onto PFNv neurons.

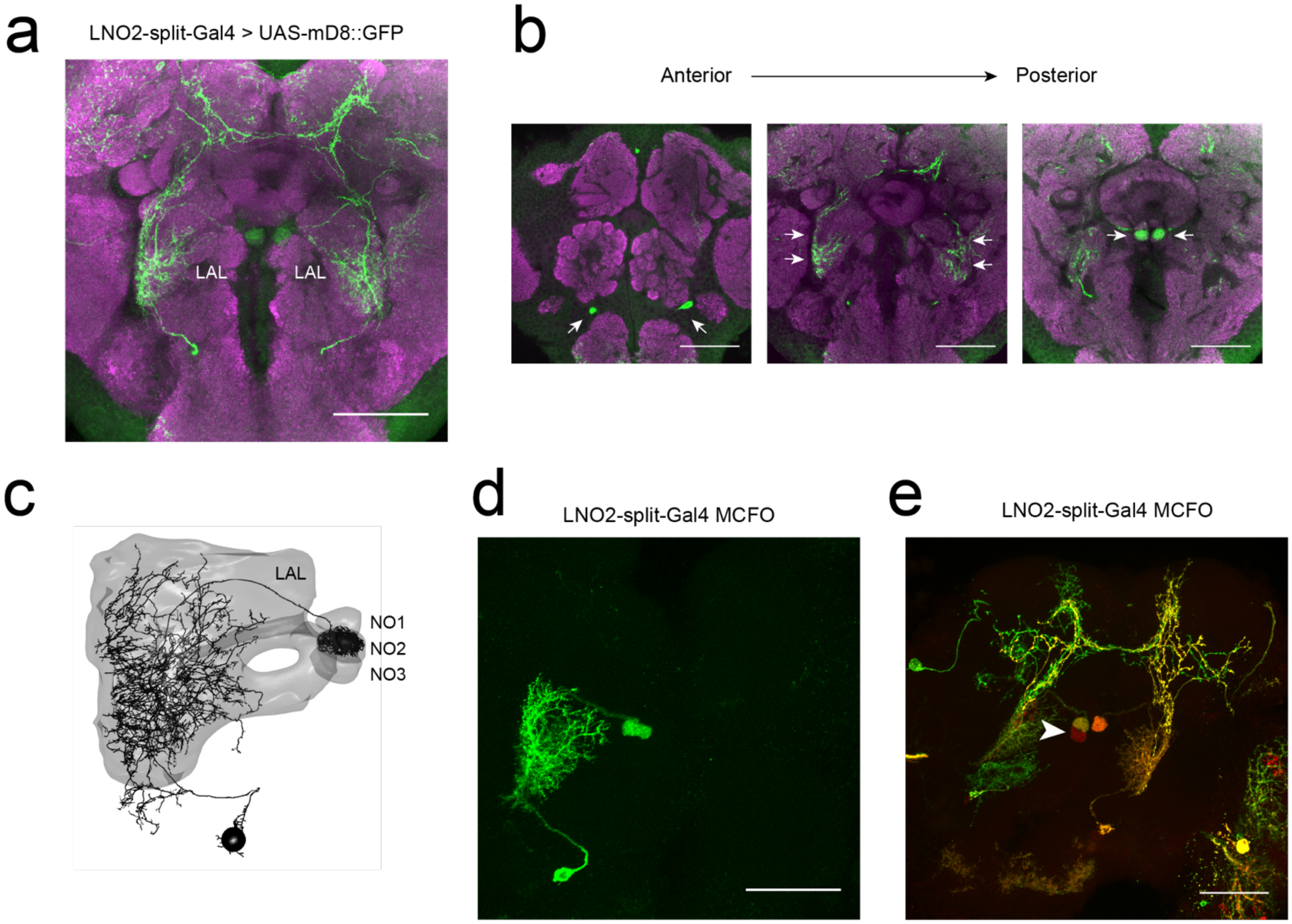


**Extended Data Figure 4: LNO2 split-Gal4 line characterization.**

a. GFP expression driven by the LNO2 split-Gal4 line: +; Mi{Trojan-p65AD.2}VGlut[MI04979-Tp65AD.2]; P{VT008681-Gal4.DBD}attP2. Shown is a coronal projection of a confocal stack through the anterior half of the brain. GFP staining is shown in green, and neuropil staining (nc82) is shown in magenta. The scale bar is 50 µm. Note that, in addition to targeting LNO2 neurons in the LAL, there are some cells are labeled in the superior brain which are not LNO2 cells.

b. Same as (a) but for individual optical slices. Shown are the location of the LNO2 cell bodies (left, arrows), neurites in the LAL (middle, arrows), and neurites in NO2 (right, arrows). Scale bars are 50 µm.

c. Skeleton of LNO2 neuron from the hemibrain dataset. Overlaid are the anatomical boundaries of the LAL and the NO (divided into subunits NO1, NO2, and NO3). The black sphere denotes the position of the cell body. There is one LNO2 neuron per hemisphere.

d. MCFO labeling of a single LNO2 neuron from the LNO2-split Gal4 line. Scale bar is 50 µm.

e. On occasion, the LNO2 split-Gal4 line shows expression in NO3. Shown is an MCFO sample from the LNO2-split Gal4 line that labels this additional neuron in NO3 (arrow). Given that two channels (green and red) label the LNO2 on the ipsilateral side, whereas only one channel (red) shows the NO3-innervating neuron, this neuron appears to be a distinct neuron from LNO2.

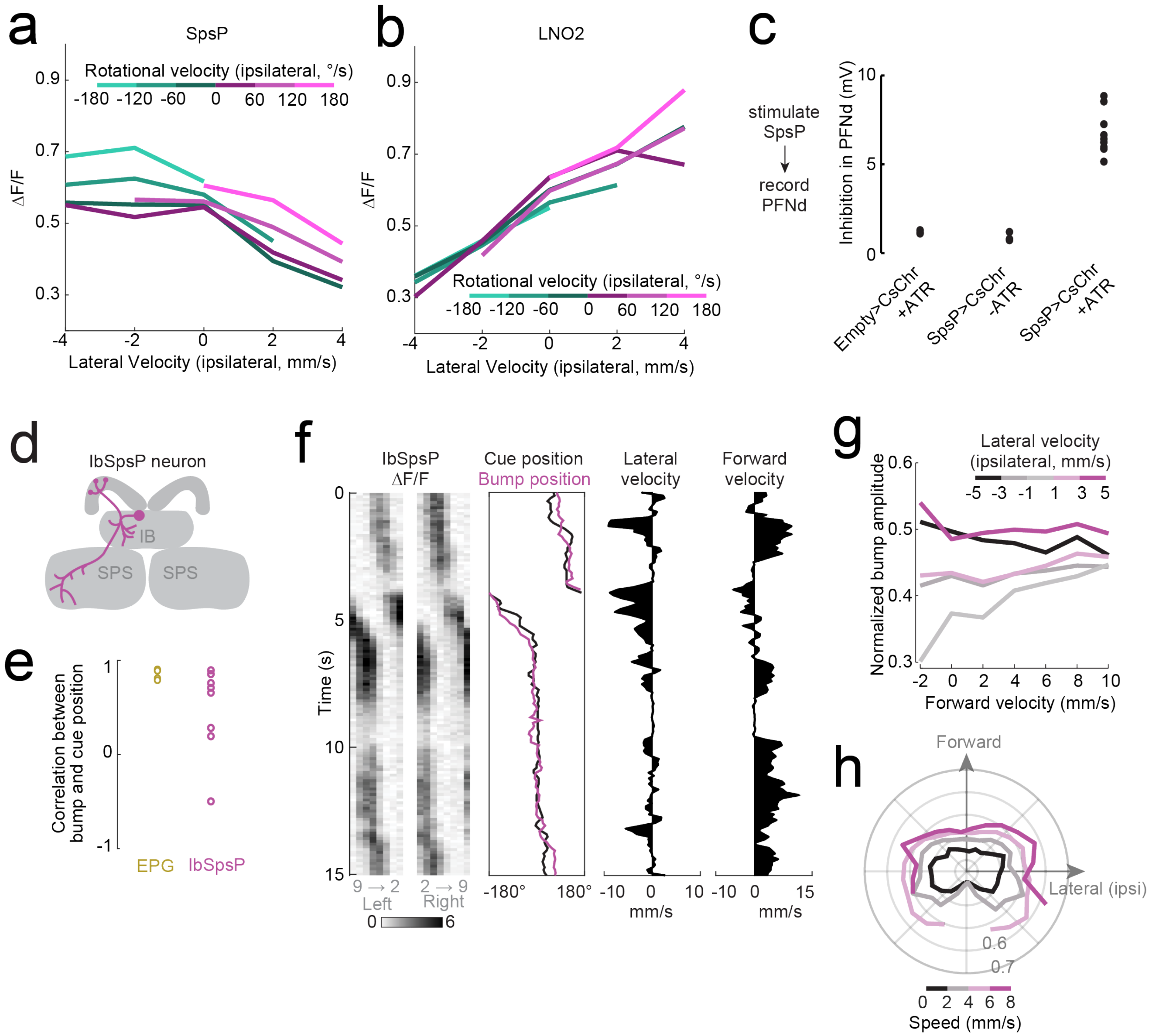


**Extended Data Figure 5: SpsP, LNO2, and IbSpsP physiology.**

a. SpsP ΔF/F versus lateral velocity in the ipsilateral direction. Data for the right and left PB are combined, binned by the ipsilateral rotational velocity, and averaged across flies (n=8 flies). Because rotational and lateral velocity is correlated, rotational velocity bins are asymmetrically populated. Note that SpsP activity increases when rotational speed is high, for both ipsi- and contralateral rotations.

b. Same as (b) but for LNO2 (n=4 flies). Note that there is little systematic effect of rotational velocity once we account for lateral velocity.

c. Control experiments for SpsP optogenetic activation. There is little effect of light in PFNd recordings from flies where an empty split-Gal4 line is combined with UAS-CsChrimson (n=3) or in flies with UAS-CsChrimson expressed under SpsP split-Gal4 control (ss52267) but reared in the absence of all-trans-retinal (ATR; n=3). We consistently see strong inhibition in flies that express UAS-CsChrimson under SpsP split-Gal4 control (ss52267) and that are raised on culture media containing ATR (n=9, reproduced from Fig. 2g). PFNd recordings were performed in TTX to isolate monosynaptic responses (see Methods).

d. Each IbSpsP neuron receives input from the inferior bridge (IB) and superior posterior slope (SPS) and projects to a few adjacent PB glomeruli.

e. Circular correlation between visual cue position and IbSpsP bump position (n=8 flies). Shown for comparison is the circular correlation for EPG neurons (n=5 flies), reproduced from Extended Data Fig. 2a.

f. IbSpsP population activity in the PB as a fly walks in closed loop with a visual cue.

g. Normalized IbSpsP bump amplitude versus forward velocity. Data are binned by lateral velocity in the ipsilateral direction, combined for the right and left PB, and averaged across flies (n=8 flies). These neurons prefer high lateral speeds in either direction. When lateral speed is low, they prefer high forward velocity.

h. Normalized IbSpsP bump amplitude in the PB, versus body-centric translational direction. Data are binned by speed. Lateral velocity is expressed in the direction ipsilateral to the imaged PB, allowing us to combine data from the right and left PB before averaging across flies (n=8 flies).

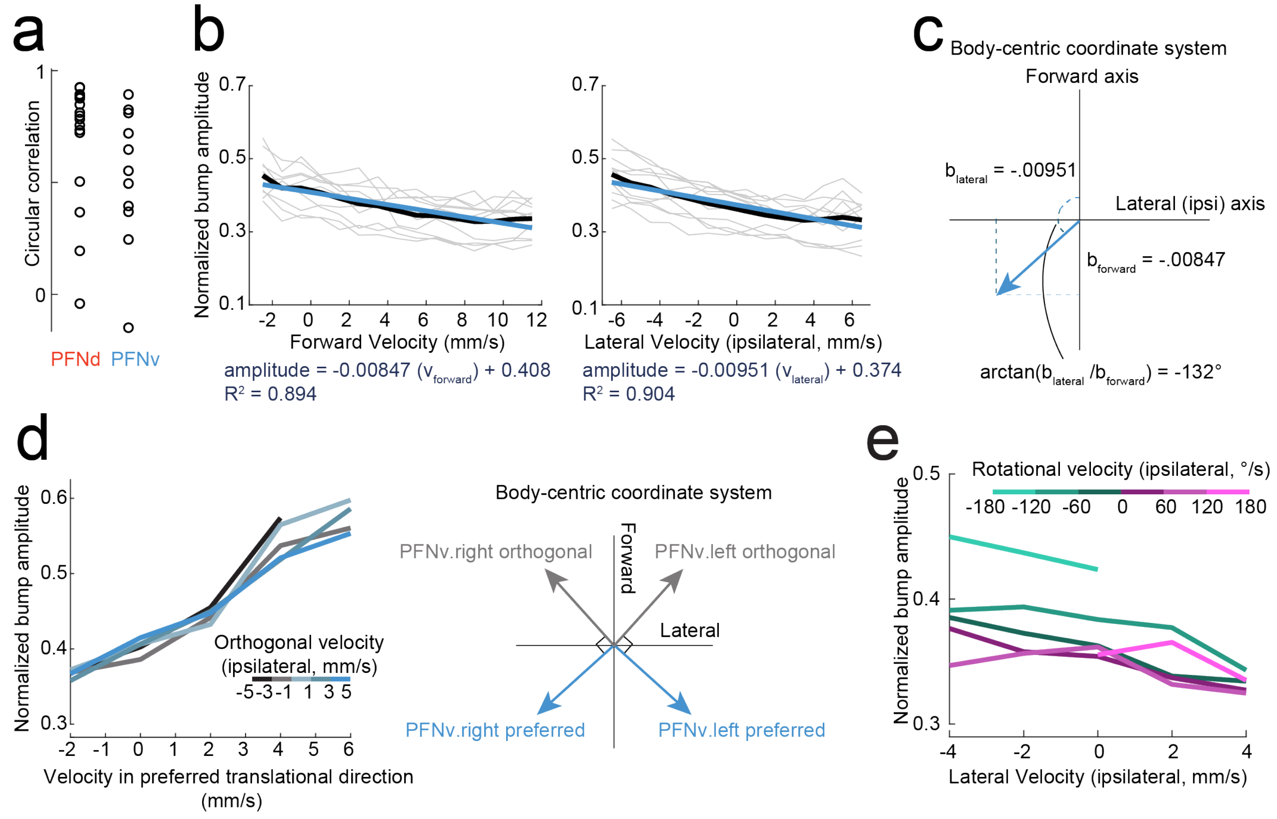


**Extended Data Figure 6: PFNv tuning properties.**

a. Circular correlation between bump and cue position for PFNd (n=16 flies, reproduced from Extended Data Fig. 2a) and PFNv neurons (n=11 flies). The measured correlation is somewhat lower for PFNv neurons, probably because the PFNv bump amplitude is often low, due to the fact that walking is generally in the forward direction.

b. Normalized PFNv PB bump amplitude versus forward velocity (left), and lateral velocity (right). Gray lines correspond to individual flies and the black line corresponds to the mean across flies (n=11 flies). Data for the right and left PB are combined. Lateral velocity is computed in the ipsilateral direction (so that, for PFNv.L neurons, leftward lateral velocity is positive and rightward lateral velocity is negative). The blue line shows the linear fit to the mean line, with the fitted equation below each plot.

c. Computation of preferred translational direction angle using the linear regression slopes for forward and lateral velocity. We used ratio of the slopes of the linear fits to lateral and forward velocity to calculate the angle of preferred translational direction. For PFNv, we found that the angle of preferred translational direction was -132° from the forward axis in the ipsilateral direction. This is equivalent to 48° from the backwards axis in the contralateral direction.

d. Normalized PFNv bump amplitude versus velocity along the angle of preferred translational direction (left). Data are combined between the right and left PB and binned by the velocity along the angle of translational movement orthogonal to the preferred direction. Shown is the mean across flies (n=11 flies). The orthogonal directions for the right and left PFNv population are shown (right); note that a positive value in the orthogonal axis remains in the contralateral direction for the given right/left population.

e. Normalized PFNv bump amplitude versus lateral velocity in the ipsilateral direction. Data for the right and left PB are combined, binned by the ipsilateral rotational velocity, and averaged across flies (n=11 flies). Because rotational and lateral velocity is correlated, rotational velocity bins are asymmetrically populated.

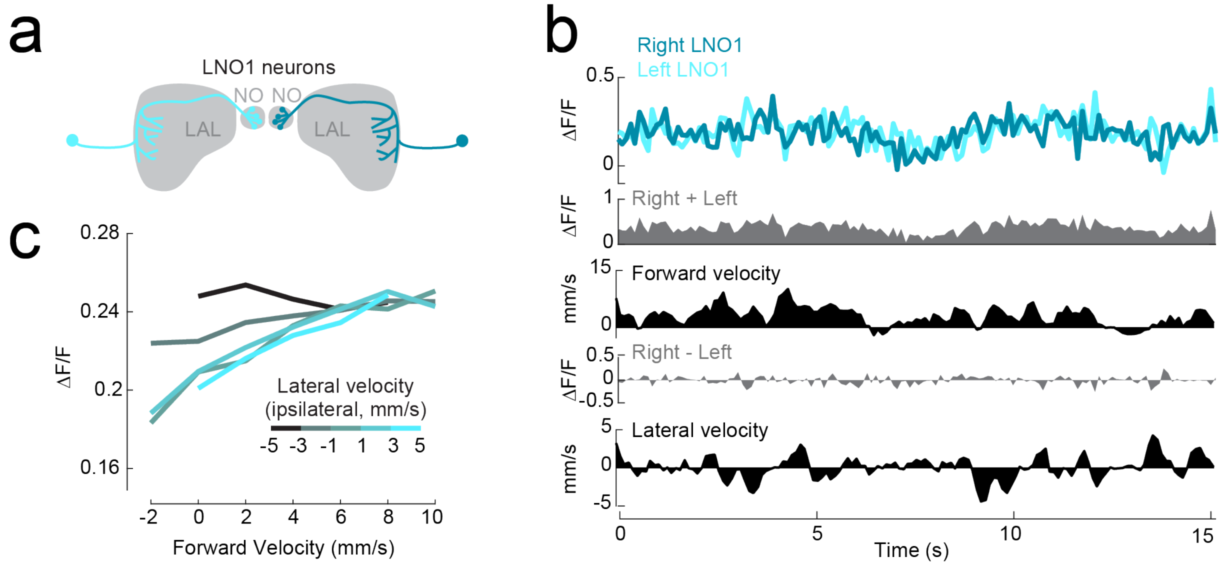


**Extended Data Figure 7: LNO1 activity versus translational velocity.**

a. Each LNO1 neuron receives input from the LAL and synapses onto PFNv and PFNd dendrites in the ipsilateral NO.

b. LNO1 activity as a fly walks in closed loop with a visual cue.

c. LNO1 activity versus forward velocity. Data for the left and right NO are combined, binned by lateral velocity in the ipsilateral direction, and averaged across flies (n=8 flies). LNO1 activity grows slightly with forward velocity, provided that the fly is not also moving laterally in the contralateral direction. Compared to the signals we observe in SpsP and LNO2 neurons, however, these signals are very small.

Note that jGCaMP7s was used in these experiments (rather than jGCaMP7f), due to the fact that LNO1 fluorescence was very dim with jGCaMP7f (see Methods).

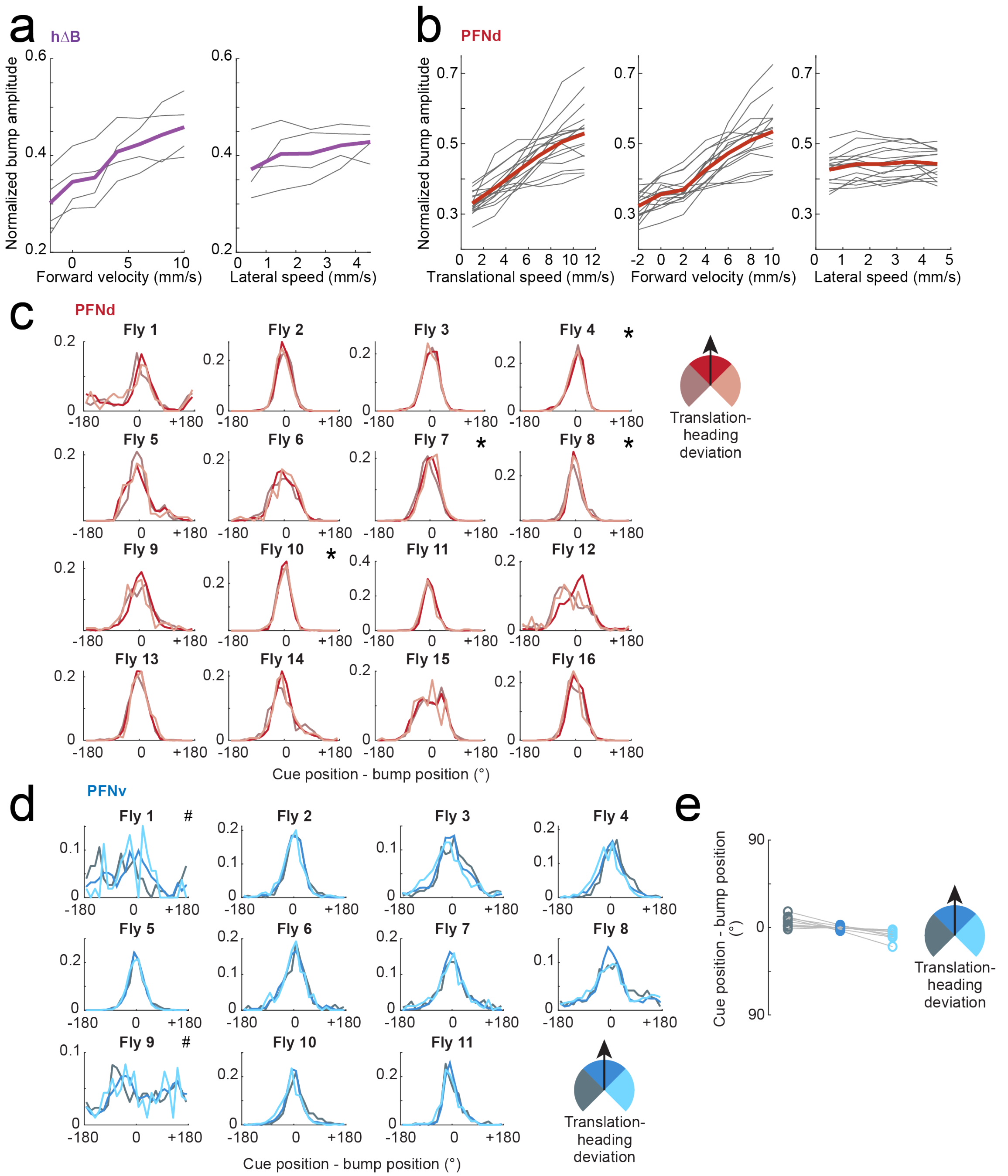


**Extended Data Figure 8: Comparing hΔB with PFNd and PFNv.**

a. hΔB normalized bump amplitude versus forward velocity and lateral speed. Gray lines show individual flies; purple is the mean across flies (n=4 flies).

b. PFNd normalized bump amplitude versus translational speed, forward velocity, and lateral speed. Gray lines show individual flies; red is the mean across flies (n=16 flies).

c. PFNd: histogram of the difference between cue position and bump position. The difference values are mean-centered within each fly, and the data is binned by the translational angle deviation (* = reproduced in Fig. 4h).

d. Same as (c) but for PFNv (# = relatively poor correlation between heading and bump position; these experiments were omitted in Extended Data Fig. 8e).

e. Mean difference between the cue position and the PFNv bump position. Each set of connected symbols is one experiment (n=9 flies; 2 flies were omitted from our analysis). Note that there is a slight shift in the opposite direction for PFNv neurons when compared to the shift observed for hΔB neurons. This shift is significant when comparing left translation-heading deviations to centered translation-heading deviations (P=0.0084, paired-sample t-test with Bonferroni-corrected α = 0.0167, Bonferroni-corrected CI = [0.0148, 0.2107] radians) and when comparing right translation-heading deviations to centered translation-heading deviations (P=0.0030, paired-sample t-test with Bonferroni-corrected α = 0.0167, Bonferroni-corrected CI = [0.0332,0.204] radians).
